# Among-Population Differentiation in the Tapeworm Proteome through Prediction of Excretory/Secretory and Transmembrane Proteins in *Schistocephalus solidus*

**DOI:** 10.1101/2024.10.25.618520

**Authors:** Anni Wang, Daniel Bolnick

**Affiliations:** Department of Ecology and Evolutionary Biology, University of Connecticut, Storrs, CT, 06269, USA

**Keywords:** Parasite, Cestode, Proteomics, Secretome, Stickleback, Coevolution

## Abstract

**Background:** Parasites secrete and excrete a variety of molecules evolve to help establish and sustain infections within hosts. Parasite adaptation to their host may lead to between-population divergence in these excretory and secretory products (ESPs), but few studies have tested for intraspecific variation in helminth proteomes.

**Methods:** *Schistocephalus solidus* is a cestode that parasitizes three spined stickleback, *Gasterosteus aculeatus*. We used an ultra-performance liquid chromatography-mass spectrometry protocol to characterize the ESP and whole-body proteome of S. solidus. Specifically, we characterized the proteome of S. solidus at the plerocercoid stage from wild caught stickleback from three lakes on Vancouver Island (British Columbia, Canada) and one lake in Alaska (United States). We tested for differences in proteome composition among the four populations and specifically between ESPs and body tissue.

**Results:** Overall, we identified 1362 proteins in the total proteome of S. solidus, with 542 of the 1362 proteins detected exclusively in the ESPs. Of the ESP proteins, we found signaling peptides and transmembrane proteins that were previously not detected or characterized in S. solidus. We also found protein spectrum counts greatly varied between all lake populations.

**Conclusions:** These population-level differences were observed in both ESP and tissue types. Our study suggests that S. solidus can excrete and secrete a wide range of proteins which are distinct among populations. These differences might reflect plastic responses to host genotype differences, or evolved adaptations by Schistocephalus to different local host populations.

## INTRODUCTION

Parasitic worms, or helminths can manipulate their hosts through various intricate strategies to establish infections which facilitates long-term survival. Among these strategies, one prominent tactic involves the production and subsequent release of an extensive range of excretory and secretory products (ESPs), which serve as pivotal mediators in the interplay of host-parasite interactions [1]. These molecules can help parasites escape the host immune system, alter host behavior, and modify host physiology [2]. Among helminth parasites, protein ESPs may include proteinase inhibitors, proteinases, heat shock proteins, and venom allergen-like proteins [3–5]. Extensive studies on helminth-derived ESP proteins or subsets of protein mixtures have demonstrated partial or total protective responses in hosts with these helminth infections [6–7]. ESPs have been analyzed in parasitic helminths such as *Strongyloides ratti*, *Ascaris suum*, *Trichuris muris*, and *Brugia malayi* [8–11].

Despite the extensive work done on helminth manipulation of host traits, some important gaps remain. For instance, intraspecific variation in helminth proteomes have been mostly overlooked, with work focusing on describing species-level proteome composition and function. This oversight is important because helminths engage in a coevolutionary arms race with their hosts [12–13]. The parasites evolve new immunosuppressive strategies while the hosts evolve new counter-measures. This coevolution should lead to rapid evolution of both species. When this rapid evolution takes place independently in geographically separate populations [14], we may expect that parasite immune modulation traits will differ among populations. This may lead to among-population divergence in the composition of the parasite proteome. Yet, intraspecific geographic variation in parasite proteomes has not been investigated. Here, we begin to rectify this research gap, by 1) describing the proteome of a widely studied tapeworm parasite and 2) testing for among-population differences in the proteome.

*Schistocephalus* is a genus of tapeworms within the family Diphyllobothriidae, specializing in parasitizing fish hosts as plerocercoid larvae through the ingestion of parasitized copepods [15]. *Schistocephalus solidus* is recognized for its infection within the threespine stickleback, *Gasterosteus aculeatus*, as it is well-documented for studying ecological processes and the genetic architecture of evolution in the wild [16–18]. *Schistocephalus solidus* exhibits a complex life cycle that requires transmission through two intermediate hosts and a final bird host. The first intermediate host is a cyclopoid copepod parasitized by the coracidium stage of *S. solidus*, which hatches from an egg deposited in the water. *S. solidus* then develops into its procercoid stage in the copepod gut, which is subsequently ingested by the second intermediate host, *G. aculeatus*. Within the fish, *S. solidus* penetrates the gut wall to enter the peritoneal cavity, where it develops into its plerocercoid stage and grows to reproductive size. If the fish is consumed by a fish-eating bird, the tapeworm matures into an adult within the host intestine, where it mates (or selves) and produces eggs. Sticklebacks infected at the plerocercoid stage have been documented to have an altered immune system, morphology, and behavior, making the plerocercoid an appealing stage to study potential ESPs responsible for these changes [19–21].

Initial proteomic work on the host-parasite system focused on characterizing subsets of proteases and transferases responsible for different developmental stages in *S. solidus* that promote growth and survival [22–23]. Subsequent proteomic work has focused on characterizing the parasite proteome and ESP at the plerocercoid stage, detection of host proteome changes due to ESPs, and parasite ESPs increasing respiratory burst activity in stickleback [24–26]. While these studies have helped establish a reference proteome and identify potential ESPs interacting with the host proteome, they have mostly not considered the possibility that the proteome evolves and thus differs among populations.

Many freshwater populations of stickleback have evolved resistance to *S. solidus* [27]. Some lake populations are more resistant, initiating an extensive peritoneal fibrosis response that suppresses tapeworm growth and viability; other lake populations evolved a tolerance strategy by suppressing fibrosis [28–29]. Transcriptomic analysis reveals among-population variation in fish immune genes in stickleback fish to *S. solidus* across different lakes [30–31], which can be induced by injecting *S. solidus* proteins into the fish [32]. However, host fibrosis response also depends on *S. solidus* genotype [33], suggesting that both host and parasite are coevolving. We hypothesized that this coevolution would lead to divergence in *S. solidus* proteome composition among plerocercoids from diverse lake populations. We further hypothesized that among-population divergence would be greater for excretory/secretory products, than for tissue proteomes, potentially influencing diverse host-parasite interactions [29]. To test these hypotheses, we conducted proteomic profiling of *S. solidus* tissue and their ESPs sampled from stickleback hosts in four distinct lakes in British Columbia, Canada and Alaska, USA. Specifically, we identified signaling peptides, excreted proteins, transmembrane proteins, and differences in proteome composition across these different lakes that could be molecules responsible for establishing infection or maintaining host-parasite interactions.

## METHODS

### Fish and Tapeworm Collection

We caught threespine stickleback from three lakes on Vancouver Island (Boot, Nimpkish, and Roselle Lakes) and one lake from Alaska (Walby Lake; geographic coordinates listed in Supplementary Table S1, sampling methods Figure 1). The four lakes chosen as field sites are associated with information from previous years in which parasites and hosts were collected to document infection rate and parasite prevalence. Roselle and Nimpkish Lake were chosen as they were geographically close to each other and we suspected that the tapeworm proteome from these two sites would be similar since genomic data show the genomes of tapeworms from these two lakes to be very similar. We chose Boot Lake as it is about 200 km away from Roselle and Nimpkish Lake and prior research conducted has shown that heavy parasitism causes fish to establish an aggressive fibrotic-like immune response (Bolnick, pers. obs). Walby was chosen as a collection site from Alaska, farthest away from the Vancouver Island lakes so we expect to see an especially distinctive proteome from Walby Lake tapeworms.

**Figure 1.**
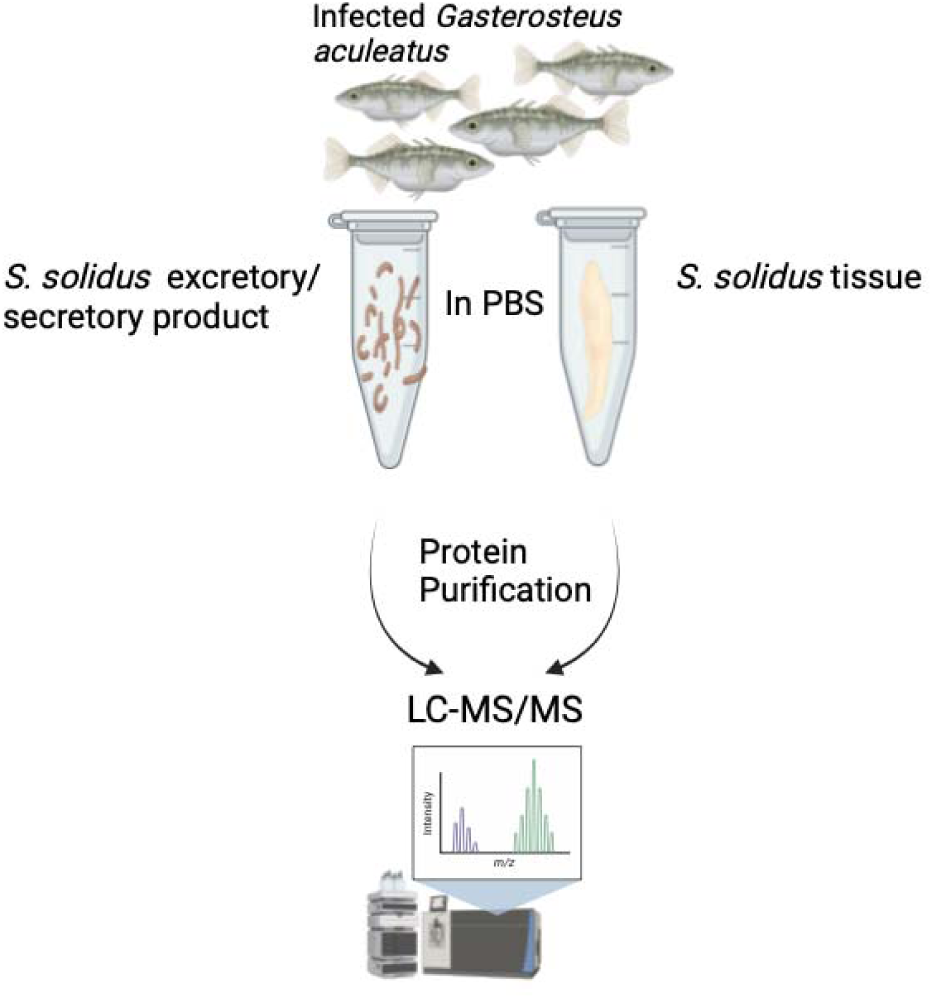
Graphical illustration of approach for separating the ESP and tissue proteins for studying the S. solidus proteome. Infected three-spine stickleback is first caught and then each *S. solidus* tapeworm is placed in PBS for two hours before ESP and tissue proteins are extracted and purified separately for LC-MS/MS analysis. Created with BioRender.com (agreement number PW270IZDQA).

**Figure 2.**
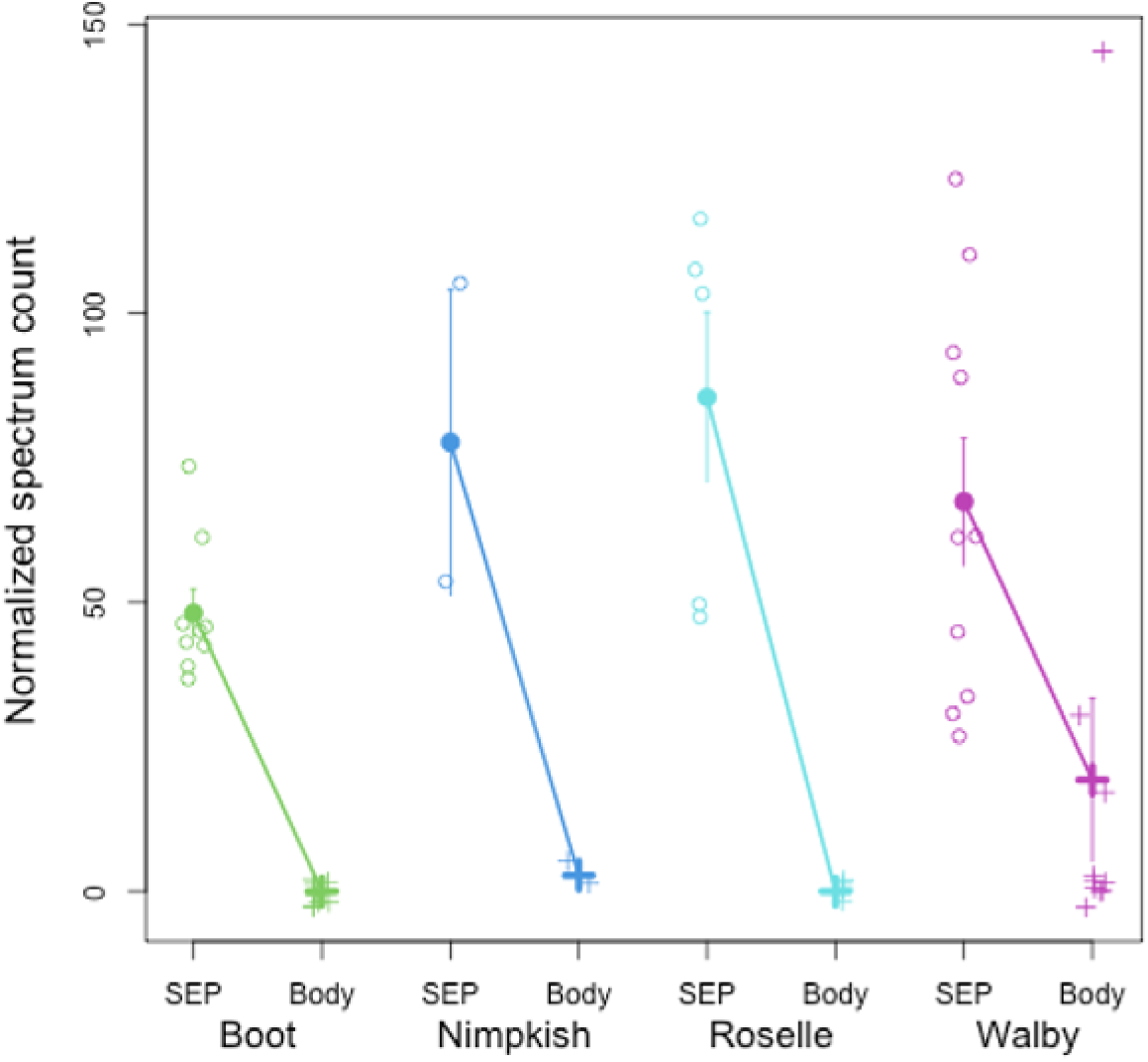
Normalized spectrum count of paramyosin. SEP stands for secretory/excretory protein and body indicates paramyosin found in the tissue proteome. Raw observations are plotted in open circles, with means and 1 standard error confidence intervals for each tissue within each population. A general linear model confirms that paramyosin exhibits statistically significant differences between ESP vs tissue (P < 0.0001), among populations (P = 0.0159), and a population*tissue interaction (P < 0.0001) indicating that the ESP-tissue difference is larger in some lakes than others.

Fish were collected in unbaited minnow traps. Sampling was conducted with the approval of the British Columbia Ministry of Environment (permit NA22-679623) and the University of Connecticut IACUC committee (protocol No. A21-025). Fish were euthanized in MS-222, chilled at 3□, and were quickly flown back to the University of Connecticut for dissection. Any live tapeworms dissected out of the fish were individually weighed and immediately rinsed with phosphate-buffer saline (PBS) before being placed in individual 2 ml tubes with 1000 ul of PBS following a protocol adopted from Berger et al. (2021). 27 tapeworms were collected, five from fish with only a single S. solidus individual and 22 from fish with multiple S. solidus individuals (Table 1). Thirty fish from each lake were dissected to find enough tapeworms, but only two were found from Nimpkish Lake fish. Each tube was covered with aluminum foil to protect each tapeworm from light exposure that could alter the proteome composition under novel environments and to mimic the dimness of the fish peritoneal cavity where plerocercoid larvae of S. solidus are typically found. A protease inhibitor was not added to preserve proteins in the secretome as the tapeworms release proteins, because previous studies have shown this induces changes to the proteome composition (Bolnick et al., 2023). After two hours in the tube, the tapeworm (for tissue) and PBS (for ESPs) were separately collected, flash frozen, and stored at -80 C for proteomic analysis.

**Table 1.**
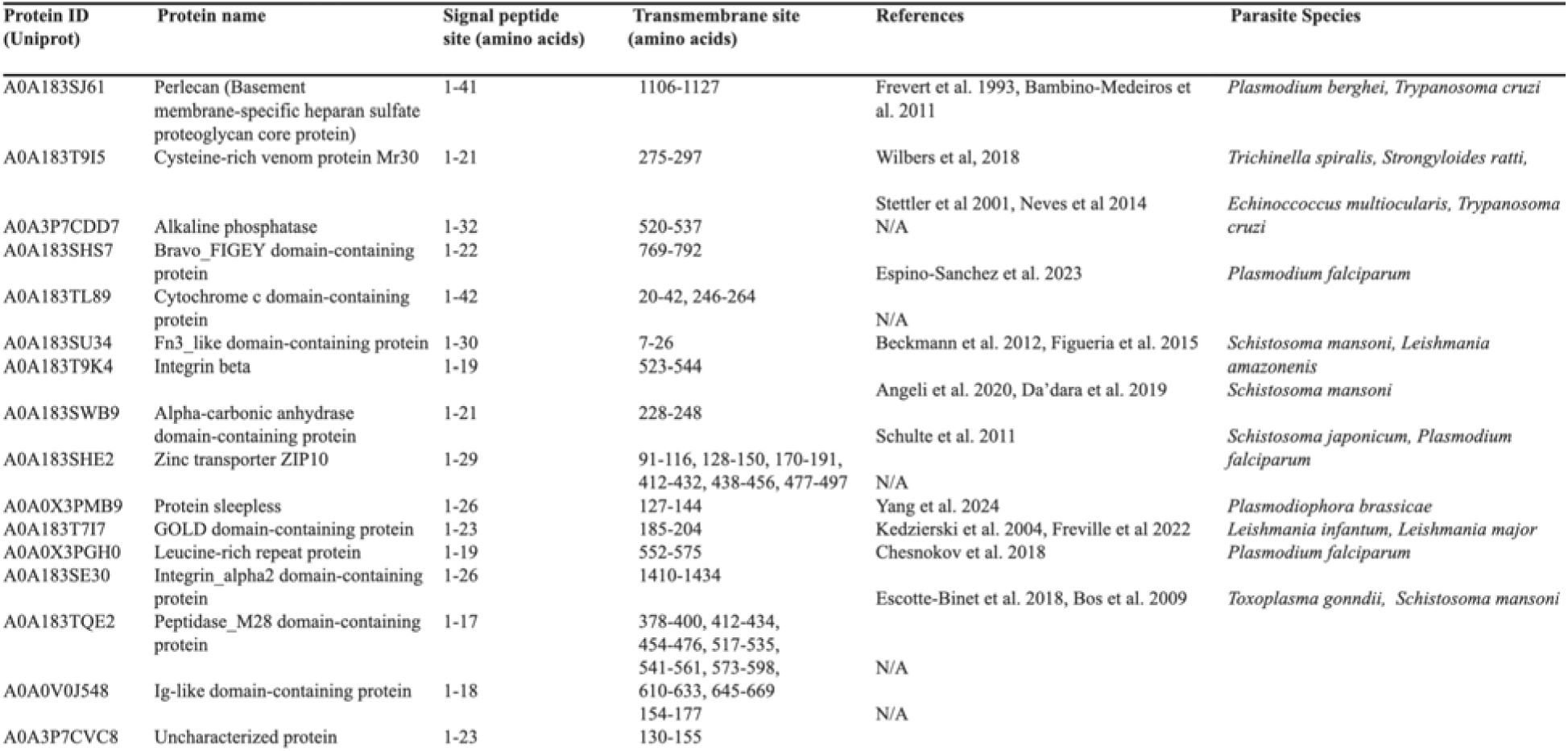
Proteins detected as signaling peptide and transmembrane bound.

For each tapeworm tissue sample, tissue was cut and placed in a tube containing lysis buffer, sodium dodecyl sulfate with Tris, and dithiothreitol. The scolex and strobilar portions of the tapeworm were both used in sample preparation. A mixture of sterile ceramic and glass beads was added to each tube and samples were homogenized for one minute at 5000 rpm using a Fisherbrand Bead Mill Homogenizer. Homogenization was repeated for three cycles, with samples placed on ice in between each cycle to prevent overheating. Samples were then centrifuged at 8,000xg for two minutes and the supernatant of each sample was transferred to new tubes. Protein concentrations were measured on a NanoDrop spectrophotometer (A280 nm) before proceeding to the protein digestion protocol. For secretome samples collected in PBS, protein concentrations were directly measured before proceeding to the same digestion protocol.

### Untargeted Protein Identification and Label-free Quantification via Tandem Mass Spectrometry In-solution digestion protocol for protein purification

For reduction/alkylation of Cys residues, 200 mM of ammonium biocarbonate was added to each sample at a 1:1 sample:buffer ratio to make a working concentration of 100mM (pH 8). 100mM of fresh dithiothreitol (6ul for each sample) was then added to yield a final sample concentration of 5mM (pH 8), and then reduced for 90 minutes at room temperature on a shaker. The sample was then alkylated in the dark with 100mM of fresh iodoacetamide (11ul for each sample) for 45 minutes at room temperature on a shaker. Once samples were alkylated, proteolysis using a modified porcine trypsin protease (Promega #V5113) was added at a ratio of 1:20 w/w enzyme:protein at 37°C and incubated overnight. The next day samples were acidified with concentrated formic acid (pH 3).

Samples were then transferred to a Pierce peptide desalting high-capacity spin column (Thermo Fisher #89852) and desalted by reverse-phase chromatography. To prepare spin columns for desalting, columns were first centrifuged at 5000 xG, then washed with pure acetonitrile twice before being finally washed with buffer A (0.1% formic acid diluted in Fisher Optima LC/MS grade water). Acidified samples were loaded into each spin column and washed twice with buffer A, eluded twice with an elution buffer (50% acetonitrile in 0.1% formic acid), and finally allowed to dry completely before resuspension in buffer A (0.1% formic acid in Fisher Optima LC/MS grade water). Samples were quantified by A280 absorbance and total concentrations were normalized.

### LC-MS analysis

*Schistocephalus solidus* samples (500 ng each) were analyzed using a Thermo Scientific Ultimate 3000 RSLCnano ultra-high performance liquid chromatography (UPLC) system coupled to a high-resolution Thermo Scientific Q Exactive HF mass spectrometer. Each sample was injected onto a PepMap RSLC C18 column (2 μm, 75 μm x 25 cm, Thermo Fisher #ES902) and separated by reversed-phase UPLC using a gradient of 4-30% Solvent B (0.1% formic acid in Fisher Optima LC/MS grade acetonitrile) over a 50-min gradient and 30-90% Solvent B over 10 min, followed by column washing and re-equilibration, at 300 nL/min flow and 50°C. Peptides were eluted directly into the Q Exactive HF using positive mode nanoflow electrospray ionization. MS1 scans were acquired at 60,000 resolution, with an AGC target of 1e6, maximum ion time of 60 ms, and a scan range of 300-1800 m/z. Data-dependent MS2 scans were acquired at 15,000 resolution, with an AGC target of 1e5, maximum ion time of 40 ms, isolation window of 2.0 m/z, loop count of 15, normalized collision energy of 27, dynamic exclusion window of 30 s, and charge exclusion “on” for all unassigned, +1, and >+8 charged species.

### Data processing

Peptides from all samples were identified using MaxQuant software (v1.6.10.43) and its embedded Andromeda search engine, and quantified using label-free quantification (Maeda et al., 2019). Raw data from the samples were searched against the UniProt S. solidus proteome (identifier UP000275846, accessed 03/27/2022), the Caenorhabditis elegans reference proteome (UP000001940, accessed 12/15/22), and the MaxQuant contaminants database. The minimum peptide length was set to five residues for the search. Variable modifications allowed oxidation of Met, acetylation of protein N-termini, deamidation of Asn/Gln, and peptide N-terminal Gln to pyroGlu conversion. Carbamidomethylation of Cys was set as a fixed modification. Protease specificity was set to trypsin/P with a maximum of two missed cleavages. All results were filtered to a 1% false discovery rate at the peptide and protein levels using the target-decoy approach; all other parameters were kept at default values. MaxQuant output files were imported into Scaffold (v5.1.2, Proteome Software, Inc.) for data visualization and subsequent analyses.

We created a more robust plerocercoid proteome dataset by not filtering out small fragments of proteins and peptides, as well as including tapeworms with mass less than 50 mg, a mass threshold used by some to define the parasite infective stage that begins to alter behavior in fish hosts [76,77].

Proteins identified with two unique peptides were considered accurately identified and any protein with one unique peptide was discarded. Protein transmembrane regions were predicted with DeepTMHMM (version 1.0.24, [34]) and Phobius [35]. Secreted proteins were predicted using SignalP 6.0 (using “Eukaryota” option, https://services.healthtech.dtu.dk/services/SignalP-6.0/[77]). Protein annotation was determined from BlastP on the UniProt database with default parameters. A matrix of total spectrum counts were exported and analyzed in R (version 4.4.0) to generate Non-metric Multidimensional Scaling (NMDS) axis scores for each sample [36]. We used a permutational ANOVA (perMANVOA) to test for between-group differences in the spectrum counts, either comparing tissue types (ESP vs whole body proteomes), comparing among populations of tapeworms, and testing for population by tissue interaction effects. To identify specific proteins underlying population and tissue differences, we iterated through each protein. For each protein we used a binomial general linear model to test whether normalized total spectrum counts varied as a function of tissue type, population, or a tissue*population interaction effect. We used a strict Bonferroni correction for multiple comparisons to determine statistical significance thresholds. All data and code required for analyses and graphics presented here are available on FigShare (https://doi.org/10.6084/m9.figshare.25718403.v1).

## Results & Discussion

A total of 1362 proteins were identified from tapeworm tissue and ESPs, with 439 uncharacterized proteins (available on FigShare). The ESP composition from all lakes consisted of 1285 proteins, with 542 of those being exclusively in the secretome. Proteins identified ranged from 651kDA to 5kDA in weight, anything lighter in molecular weight was not detected. Of the 1362 total proteins, 60 were predicted to have at least one signaling peptide region, 219 were predicted to have at least one transmembrane region, and 16 had both a signaling and transmembrane region (Table 2). Signaling peptides are responsible for directing proteins to their appropriate cellular location which can include functions like secretion out of a cell or targeting specific organelles within a cell. Transmembrane proteins, however, are components of cellular membranes and perform functions such as cell signaling, cell-cell communication, or transportation of molecules across membranes. Within the context of parasite proteins, the presence of both types of proteins can provide valuable insights into localization and function within a host organism. For example, detection of parasite signal peptides could indicate proteins that are secreted to play roles in modulating host immune responses or host cell functions. Parasite transmembrane proteins could interact with host cell membranes, which may lead to evasion of host defense mechanisms or facilitating parasite entry into host cells.

**Table 2.**
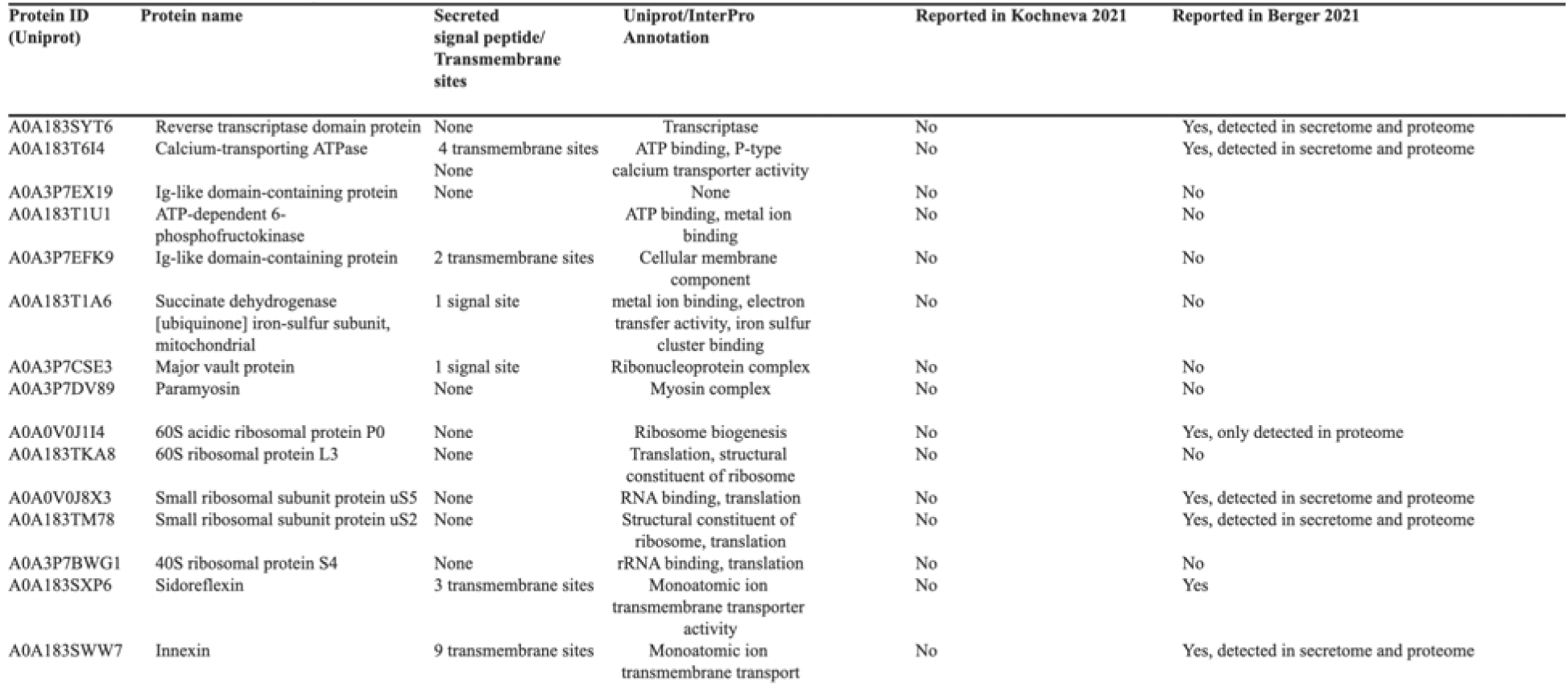
Excreted proteins detected majority in secretome than proteome. Minimum at least 2 unique peptides and spectrum count per sample, any protein with ambiguous a, b, and y ion relative intensity values were excluded, uncharacterized protein were excluded.

### Secreted signal peptides

We identified 15 proteins that were found in the secretome from all four lakes. Eight had not previously been identified in *S. solidus* literature. Two of the newly detected secretome proteins (A0A3P7EX19 and A0A3P7EFK9) resemble immunoglobulin domain proteins which may have immunoregulatory properties in regulating stickleback infections, but the proteome is not annotated well enough to suggest what those may be. Additionally, 5 of these secretory proteins belong to either the large (60S) or small (40S) ribosomal subunit family (A0A0V0J1I4, A0A183TKA8, A0A0V0J8X3, A0A183TM78, A0A3P7BWG1), responsible for ribosome synthesis and rRNA processing. We also identified paramyosin (A0A3P7DV89), a myofibrillar protein that has been shown in other helminth parasites to play a multifunctional role in host-parasite interactions [37–38]. Across all lake populations, paramyosin had an 8.8-fold higher total spectrum count in the ESP than in the tissue (P < 0.0001). Paramyosin is a rod shaped α-helical myosin-binding protein commonly found in invertebrates through linking with actin filaments and tropomyosin, most likely involved in specialized contractile functions [39]. Paramyosin in *Clonorchis sinesis*, *Schistosoma mansoni*, and *Trichinella spiralis* have been shown to bind and stimulate host complement component proteins, which allows the parasite to evade the host’s innate immune system [37–38, 40].

### Secreted and Transmembrane Proteins

Within the 16 identified proteins that had both a signaling peptide and transmembrane region, 11 were found in other parasites with known functions of maintaining host-parasite interactions and five in helminth parasites (Table 2). One protein (A0A183T9I5) is a secreted venom allergen like (VAL) protein that is structurally conserved and abundantly secreted in multiple stages of helminth parasites [41]. VALs were initially studied for vaccine development due to their protective immunity in parasites as they aid in transmission, tissue migration, and feeding. In animals, parasites produce VALs from secretory glands and modulate host immune responses through inhibition of platelet aggregation, granulocyte adhesion, oxidative burst, and B cell signaling pathways [42–44].

### Proteome divergence among tissues and among populations

To compare the ESP and whole tissue proteome, we performed an NMDS with K=8 to differentiate all eight groups (four lake populations by two sample types). Loadings of proteins on the first four NMDS axes are provided in Supplementary Table S3. A multiple MANOVA applied to NMDS axes 1-4 confirmed there is a significant effect of both tissue type (Pillai’s trace = 0.87, F_2,50_ = 172.42,*P* < 0.0001), genotype of parasite (Pillai’s trace = .908, F_12,123_ = 4.45, 132, *P* < 0.0001), and a genotype by tissue interaction (Pillai’s trace = 0.674, F_12,132_ = 2.97, *P* = 0.0011). The significant genotype by tissue interaction indicates that the difference between the proteins in the body and ESP depends on the parasite population. The first NMDS axis (Fig. 3) separates ESP versus tissue proteome for all four populations. Both the first and second axes separate the populations but the third and fourth NMDS axes exhibit population-specific differences between ESP and tissue. These population-specific differences are strong candidates for proteins involved in local adaptation to their local host populations. We also observed statistically significant differences only within the ESP proteome between the different tapeworm populations (Pillai’s trace = 1.19, F_12,63_= 3.44, df = 12,63, *P* = 0.0006; Supplementary Table S4). Similarly, we observed statistically significant differences in the tapeworm tissue proteome composition between tapeworm populations (Pillai’s trace = 1.68, F_12,66_ = 6.58, *P* < 0.0001, Supplementary Table S4). Among-population variance in ESP composition is not significantly greater than among-population variance in whole body proteomes (F-test of the ratio of variances, F_4,4_=1.285, P=0.2830). This suggests that both ESP and tissue proteomes are diverging among populations, and the ESP proteins are not uniquely important for population adaptation or coevolution.

**Figure 3.**
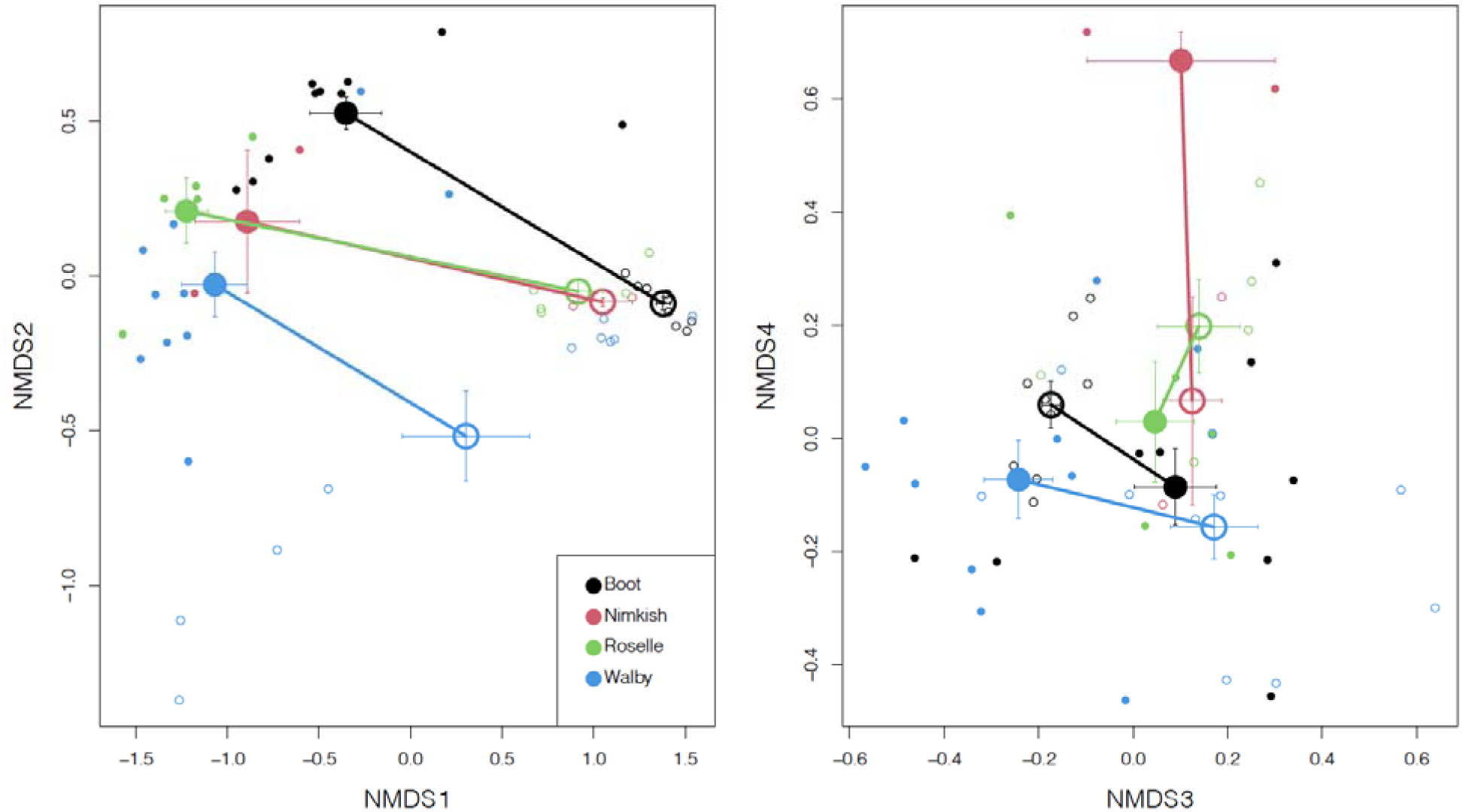
Lake origin and differences in the ESP and tissue proteome were examined, lake origin affects differences in proteome more than ESP vs. tissue proteins. We used the elusive spectrum count from each protein found in each fish to calculate NMDS scores. Lake origin is colored coded with larger circle and symbol to denote standard error bars. 2a shows NMDS of ESP and tissue proteome, with ESP in black and tissue proteome in red. 3b shows the entire proteome combined for all lakes. The closed and open circles represent tissue and ESP proteins respectively.

Analyses of each protein separately also identified many that significantly differed between tissues, and between populations (Supplementary Table S5). We found a cluster of six actin proteins (A0A183SVY0, A0A183T1Z7, A0A0X3PDD0, A0A183TJY3, A0A183S955, A0A183TGM4) that were significantly different in the ESP and tapeworm tissue (all *P* < 0.0001), as well as significant difference in the level of actin abundance within tapeworm tissue among the different lakes. Two paramyosin proteins (A0A183T4B5, A0A3P7CMF5) appear to be differentially abundant across different lakes (*P*<0.0001) as well as between the ESP and tissue proteome (*P*<0.0001). Tapeworm tissue either did not contain either proteins or had a spectrum count of less than 10. Both proteins have spectrum counts above 100 in the ESP of all tapeworms except for six from Walby Lake, with total spectra highest in Roselle Lake ESP. Other muscle proteins involved in muscle formation like transgelin (A0A183SYR3), tubulin (A0A183TBY5), and myosin (A0A183T9Q5) were found to have higher spectrum counts in the ESP than tissue proteome (all *P*<0.0001), with the tissue proteome from Nimpkish, Roselle, and Walby Lake typically show two spectrum counts or less. Immunohistochemical studies in other tapeworms like *E. granulosus* and *T. solium* have shown these myofibrillar proteins to bind to host Fc fragments on immunoglobulins, complement C1, and complement C9, making them attractive targets for parasite diagnostic tests and pharmacological development [45].

Aside from muscle proteins, we additionally identified two heat shock proteins, heat shock protein 70 (HSP70, protein ID:A0A183S9K7) and heat shock protein 90 (HSP90, protein ID:A0A183SJF6) with total spectrum counts highest in the ESP of tapeworms found in Boot Lake, but lowest in ESP of tapeworms in Walby Lake (tissue difference P < 0.0001, population effect P = 0.0003, and tissue*population interaction effect P = 0.0008). This is representative of several proteins whose body/ESP difference varies significantly among lakes (significant population*tissue interactions, Supplementary Table S4). HSPs are a family of highly conserved molecular chaperones found in eukaryotic organisms and are important in cellular processes, helping denatured proteins to refold or to target them for degradation [46]. HSPs are expressed under normal conditions but expression increases under stresses such as temperature changes, heavy metal exposure, and parasite infections. HSPs are grouped based on their molecular weight so major groups are typically denoted under HSP100, HSP90,HSP70, HSP60,HSP40, and smaller HSPs. During parasite development, transmission through different hosts during developmental stages like passing through cold-blooded hosts e.g., fish, mosquitos, insects and switching to homeothermic hosts, requires HSPs to regulate for the change in such drastically different environments.

HSPs and their role in modulating helminth’s response to their environments have been reported in several species [47–48]. HSP70 in *Echinococcus multilocularis* is found to be released as extracellular vesicles by the protoscoleces which could potentially promote angiogenesis [49]. The C-terminal region of HSP70 in *Echinococcus ganuulosus* has been shown to induce host IgG and IgE response [47]. *Chonorchis sinesis* produces both HSP70 and HSP90 in its adult stage with both proteins inducing a TH1 response in hosts [50]. HSP90 has been found in extracellular vesicles in various helminths that can regulate host macrophage functions and modulate other immune responses [47,51].

### Excreted Proteins

In a prior study of *S.solidus* from Quebec, eight proteins were reported to be found exclusively in the secretome of all sampled individuals [24]. We did not find any of these eight proteins in our samples. In the Berger 2021 proteome study, an additional twenty-two exclusive secretome proteins were detected only in some of the proteome samples. However, only three of these proteins (A0A183TPG4, A0A183TIR8, A0A183TE24) were identified in our dataset, none of them were exclusively found in the secretome. We believe this observation is due to differences in geographical location as well as how the threespine stickleback in the Berger et al. (2021) were raised prior to each experiment. A0A183TPG4 was only found in five tapeworms from Boot Lake and one from Nimpkish Lake. However, all five tapeworms from Boot Lake had only one unique peptide, which falls below our filtering threshold. The protein contains a cystatin domain, which may have host immunomodulatory properties. Cystatin is a protease inhibitor primarily secreted by parasites to evade host immune responses [52]. A0A183TIR8 was found in at least one tapeworm from each lake. The protein functions as a sodium/glucose cotransporter protein involved in mediating sodium and glucose transport across cell membranes. Since *S. solidus* does not possess a digestive system and instead obtains nutrients by consuming carbohydrates (or glucose) through the glycolytic pathway, proteins that regulate glycolytic processes are especially important for survival. A0A183TE24 is an intraflagellar transport protein required for the maintenance and formation of cilia. We found this protein in at least one tapeworm from every lake (11 total) in both tissue and secretory samples. However, all samples except one tapeworm from Nimpkish Lake had only one spectrum and peptide count, falling below our filtering threshold. The coracidia of *S. solidus* emerge from their eggs with cilia that aids in swimming. However, once ingested by a copepod, they shed their cilia this this peptide should not be present once they are in their second intermediate host [53].

Within the ESP, we found several excreted proteins that has been detected in some plerocercoids of *S. solidus* tapeworms found in freshwater lakes across Russia [25]. Malate dehydrogenase (A0A183TNV9) was excreted in all tapeworms, although it had previously been reported to be found only in some *S. solidus* tapeworms. We believe the discrepancy was due to varying methodology related to sample preparation for mass spectrometry. Malate dehydrogenase is an enzyme that catalyzes NAD/NADH dependent interconversion of malate and oxaloacetate, a crucial mediator in gluconeogenesis, the citric acid cycle, and as a malate-aspartate shuttle. In helminths, this protein plays an important role in obtaining energy from glucose and overcoming oxidative stress. It has been highlighted as a potential therapeutic drug target against parasites [54].

Peptidyl-prolyl cis-trans isomerase (A0A0X3Q4H0), also known as immunophilin, was found to be excreted in all tapeworms except three plerocercoids from Walby Lake. Immunophilins are highly conserved proteins that catalyze protein folding by accelerating the cis/trans isomerization of peptide bonds and are thought to have roles in chaperone activity. In parasites, immunophilins have been of interest due to evidence that they are involved in pathogenesis of infections and having strong inhibitory effects on certain parasites in culture and animal model infections, making them potential drug targets or mediators of drug action against parasites [55]. We also found a collagen-like protein (A0A183SRL7) that was excreted from all tapeworms except six plerocercoids from Walby Lake. It has been previously thought that *S. solidus* releases collagen into their hosts, however the function is unclear. It could be a waste product excreted passively or potentially stimulates fibrinogenesis in hosts [24,25]. A cluster of four annexin proteins (A0A183TKJ8, A0A183SWE1, A0A183SGJ0, A0A183S9W9) were found to be excreted in all four lake populations (Figure 4). Annexins are a family of protein with diverse functions found in parasite structure and secretions that have recognized roles in the pathogenesis of parasite infections [56]. In parasites, annexin have been found to downregulate immune response in their host, to calcify parasite cysts, in tissue development, and to modulate inflammation [58]. Due to their multifaceted functions in pathological processes, annexins have been suggested as potential therapeutic treatment against *Leishmania, T. solium, E. ganulosus, Taxoplasma gondii, Trypanosoma brucei*, and others [57,58].

**Figure 4.**
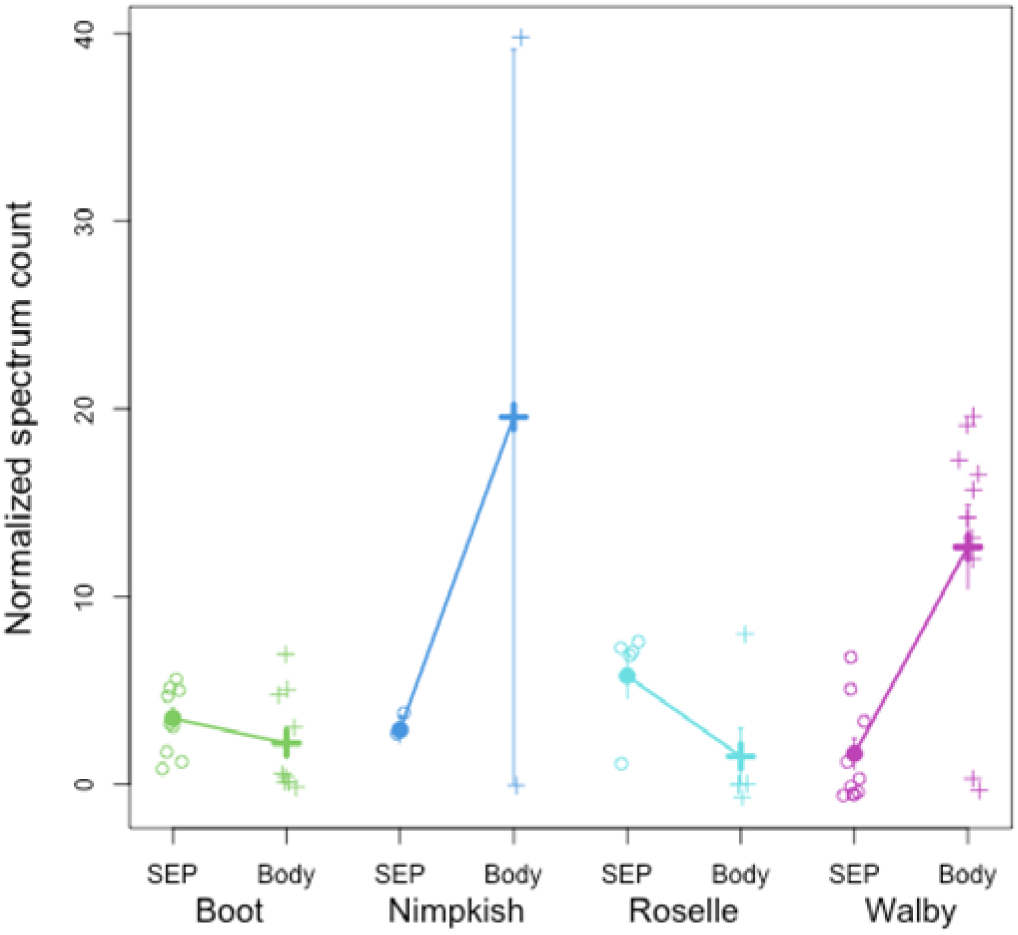
Normalized spectrum count of annexin, which shows a population by tissue interaction effect. Raw observations are plotted in open circles, with means and 1 standard error confidence intervals for each tissue within each population. A general linear model confirms that annexin exhibits statistically significant differences between SEP vs whole body (P = 0.0160), among populations (P = 0.0190), and a population*tissue interaction (P < 0.0001) indicating that the SEP-body difference is larger in some lakes than others or reverses direction.

### Lake population differences

We identified a total of 186 proteins whose spectrum counts differed significantly among the four lakes, using a strict Bonferroni false discovery rate correction. Of these 186 proteins, 60 were uncharacterized. Of the characterized proteins, 67 can be assigned to 21 protein families. Tubulin was the largest family with seven tubulin alpha chain proteins (A0A183SBQ0, A0A183SL33, A0A183T3L3, A0A183T3T2, A0A183TBY5, A0A183TD95,A0A183TJU1) and six tubulin beta chain proteins (A0A183SFW7, A0A3P7DQX4, A0A3P7D6K9, A0A3P7D9Q4, A0A0X3NM14, A0A0X3PFM5). Tubulin alpha proteins consistently had higher spectrum counts in the tissue of tapeworms compared to their ESP in three lakes (Boot, Roselle, and Walby, all P<0.0001). However, in Nimpkish Lake this protein was enriched in the ESP compared to tissue (*P*<0.0001). Tubulin beta chain protein has higher spectrum counts in the ESP than in tapeworm tissue in Nimpkish and Walby Lake only (both *P*<0.0001). Spectrum counts were higher in tubulin beta compared to tubulin alpha because beta sheets typically contain 42% of tubulin beta and 39% of tubulin alpha [59]. Tubulin alpha chain proteins are almost always detyrosinated, which makes them susceptible to faster degradation than their tyrosinated counterparts.

The second largest family that differed among lake populations was dynein, with seven dynein light chain proteins (A0A0V0JBQ8, A0A183SE96, A0A183SE97, A0A183SKR1, A0A183TQI2, A0A183TQX4, A0A0X3P671). Of interest, A0A183SKR1 had the highest spectrum count only in the tissue of tapeworms from Boot Lake (*P*<0.0001), it was nearly absent in all ESPs and tissues from all other lakes. Additionally, three paramyosin proteins (A0A3P7BUS1, A0A183T4B5, A0A3P7CMF5) exhibited higher spectrum counts (*P*<0.0001) in the ESP of all tapeworms than in tissue samples. Specifically, A0A3P7BUS1 averaged less than 10 spectrum counts, while the others had at least 50 counts or higher, with an average around 100 counts.

## Conclusions

In summary, we identified secreted, excreted, and transmembrane proteins crucial for host-parasite interactions in *S. solidus*. Highlighted proteins in the ESP may mediate these interactions, alongside new ESPs found in the *S. solidus* proteome. Comparison of ESPs and tissue proteins from different lake populations revealed previously undescribed differential expression levels in *S. solidus* proteome studies. These findings underscore the potential for microevolutionary divergence in parasite proteomes among populations within close proximity. Overall, this proteomic resource enhances understanding of host-parasite interactions and aids in identifying potential protein targets, facilitating studies of diverse interactions in nature.

## Acknowledgements

We thank Andrea Roth, Lindsay Yue, Lauren Simonse, Maria Rodgers, and Arshad Padhair for field work collection, and Swapna Subramanian for help with tapeworm dissections. We thank Jeremy Balsbaugh and Jen Liddle from the University of Connecticut Proteomics Core Facility for their advice on preparing and running samples for mass spectrometry analysis.

## Availability of data and materials

Mass spectrometry dataset generated is available on figshare, https://doi.org/10.6084/m9.figshare.25718403.v1

## Authors’ contributions

Study conception and design were done by AW and DIB. Material preparations, data collection, mass spectrometry, data analysis and first draft of the manuscript was conducted by AW. Statistical analysis was carried out by DIB. The manuscript was revised by both authors. All authors read and approved the final manuscript.

## Ethics approval

Animal experiments in this study were performed under applicable laws and guidelines for care and use of research animals, as specified in the Institutional Animal Care and Use Committee No. A21-025.

## Consent for publication

Not applicable.

