## Supplementary Tables 1-4 for "Among-Population Differentiation in the Tapeworm Proteome through Prediction of Excretory/Secretory and Transmembrane Proteins in *Schistocephalus solidus*"

Supplementary Table S1: stickleback collection site, locality, and date:

| **Lake** | **Coordinate** | **Collection date** |
| --- | --- | --- |
| Boot Lake | 50°3’15”N, 125°31’45”W | 6/16/22 |
| Roselle Lake | 50°31’13”N, 126°59’12”W | 6/16/22 |
| Nimpkish Lake | 50°32'14"N, 127°00'40"W | 6/16/22 |
| Walby Lake | 61°37'7.464"N, 149°12'53.611"W | 6/16/22 |

Supplementary Table S2: worm ID, mass, lake origin.

| **Worm ID** | **Lake Origin** | **Number of tapeworms in host** | **Mass, mg** |
| --- | --- | --- | --- |
| 46C | Boot Lake | 6 | 259 |
| 46D | Boot Lake | 6 | 268 |
| 46E | Boot Lake | 6 | 263 |
| 46F | Boot Lake | 6 | 259 |
| 53A | Boot Lake | 1 | 282 |
| 54A | Boot Lake | 1 | 298 |
| 80D | Boot Lake | 7 | 31.7 |
| 80E | Boot Lake | 7 | 71.8 |
| 80F | Boot Lake | 7 | 84.8 |
| 80G | Boot Lake | 7 | 12.8 |
| 38A | Nimpkish Lake | 1 | 51.4 |
| 40A | Nimpkish Lake | 1 | 63.5 |
| 47A | Roselle Lake | 4 | 14.9 |
| 47B | Roselle Lake | 4 | 134 |
| 47C | Roselle Lake | 4 | 116 |
| 47D | Roselle Lake | 4 | 147 |
| 48A | Roselle Lake | 1 | 207 |
| 24B | Walby Lake | 4 | 10 |
| 41B | Walby Lake | 13 | 60 |
| 41C | Walby Lake | 13 | 57 |
| 41D | Walby Lake | 13 | 67 |
| 41G | Walby Lake | 13 | 61 |
| 41K | Walby Lake | 13 | 63 |
| 41L | Walby Lake | 13 | 68 |
| 42A | Walby Lake | 2 | 73 |
| 42B | Walby Lake | 2 | 62 |
| 44B | Walby Lake | 2 | 59 |

Supplementary Table S3: NMDs loadings.

| **Protein ID** | **MDS1** | **MDS2** | **MDS3** | **MDS4** |
| --- | --- | --- | --- | --- |
| Paramyosin OS=Schistocephalus solidus OX=70667 GN=SSLN_LOCUS11313 PE=3 SV=1 | -0.3169313 | -0.9711296 | 0.23275159 | -0.1374592 |
| Paramyosin OS=Schistocephalus solidus OX=70667 GN=SSLN_LOCUS12377 PE=3 SV=1 | -0.2868979 | -1.030294 | 0.18296212 | -0.1790479 |
| Paramyosin OS=Schistocephalus solidus OX=70667 GN=SSLN_LOCUS4893 PE=3 SV=1 | -0.4703271 | -1.0387548 | 0.24082163 | -0.0908521 |
| Uncharacterized protein OS=Schistocephalus solidus OX=70667 GN=SSLN_LOCUS10987 PE=4 SV=1 | 0.0977188 | 0.29757811 | 0.2303711 | -0.634592 |
| Calponin OS=Schistocephalus solidus OX=70667 GN=SSLN_LOCUS15559 PE=3 SV=1 | -0.0482209 | 0.09233252 | 0.30552213 | -0.5426395 |
| Uncharacterized protein OS=Schistocephalus solidus OX=70667 GN=SSLN_LOCUS2612 PE=3 SV=1 | -0.3698807 | -0.9979385 | -0.005978 | 0.01927413 |
| Transgelin OS=Schistocephalus solidus OX=70667 GN=SSLN_LOCUS9361 PE=3 SV=1 | -0.3100758 | 0.13200165 | 0.03387313 | -1.0125627 |
| Major vault protein OS=Schistocephalus solidus OX=70667 GN=SSLN_LOCUS9700 PE=4 SV=1 | -0.6430337 | -1.3731515 | 0.14121058 | -0.0708064 |
| Fibrillar collagen NC1 domain-containing protein OS=Schistocephalus solidus OX=70667 GN=SSLN_LOCUS8597 PE=4 SV=1 | 0.14831471 | -0.1719734 | 0.14382301 | -0.5629471 |
| Major vault protein OS=Schistocephalus solidus OX=70667 GN=SSLN_LOCUS14029 PE=4 SV=1 | -0.6828439 | -1.3903578 | 0.14860862 | -0.0809333 |
| Uncharacterized protein OS=Schistocephalus solidus OX=70667 GN=SSLN_LOCUS16894 PE=3 SV=1 | -0.3735468 | -1.3908595 | 0.20001182 | -0.0109381 |
| Uncharacterized protein OS=Schistocephalus solidus OX=70667 GN=SSLN_LOCUS7555 PE=4 SV=1 | -0.7133376 | -1.0327432 | -0.054008 | -0.2194714 |
| Transforming growth factor beta-1-induced transcript 1 protein OS=Schistocephalus solidus OX=70667 GN=TGFI1 PE=4 SV=1 | -0.0969301 | -0.1693073 | 0.40191396 | -0.6091943 |
| CCT-theta OS=Schistocephalus solidus OX=70667 GN=SSLN_LOCUS5678 PE=3 SV=1 | -0.7369972 | -1.5535321 | 0.03910374 | -0.0125878 |
| Peptidyl-prolyl cis-trans isomerase OS=Schistocephalus solidus OX=70667 GN=PPIA PE=3 SV=1 | -0.0865466 | 0.01466934 | 0.11184299 | -0.5217269 |
| Pyruvate dehydrogenase E1 component subunit beta OS=Schistocephalus solidus OX=70667 GN=SSLN_LOCUS5537 PE=4 SV=1 | -0.56725 | -1.1434179 | 0.14472567 | -0.0293453 |
| H15 domain-containing protein OS=Schistocephalus solidus OX=70667 GN=SSLN_LOCUS8960 PE=3 SV=1 | -0.7442291 | -1.3264898 | 0.13810765 | -0.0331218 |
| Cation_ATPase_N domain-containing protein OS=Schistocephalus solidus OX=70667 GN=SSLN_LOCUS10076 PE=4 SV=1 | -0.7008075 | -0.9567275 | 0.27066869 | 0.00691736 |
| Uncharacterized protein OS=Schistocephalus solidus OX=70667 GN=SSLN_LOCUS16091 PE=3 SV=1 | -0.5006811 | -1.2352155 | 0.44300141 | 0.20798618 |
| Phosphate carrier protein; mitochondrial OS=Schistocephalus solidus OX=70667 GN=SSLN_LOCUS17082 PE=3 SV=1 | 0.34698034 | -0.744065 | 0.75518444 | 0.20507083 |
| Uncharacterized protein OS=Schistocephalus solidus OX=70667 GN=SSLN_LOCUS10145 PE=4 SV=1 | 0.24830986 | -0.8561769 | 0.81346458 | 0.33887692 |
| 40S ribosomal protein S14a OS=Schistocephalus solidus OX=70667 GN=RS14A PE=3 SV=1 | -0.957091 | -1.5067921 | 0.04747368 | 0.16228591 |
| Uncharacterized protein OS=Schistocephalus solidus OX=70667 GN=SSLN_LOCUS18753 PE=3 SV=1 | 0.89244181 | 0.04657419 | 0.2291879 | -0.6085806 |
| SERPIN domain-containing protein OS=Schistocephalus solidus OX=70667 GN=SSLN_LOCUS10184 PE=3 SV=1 | 0.91198517 | 0.05735825 | 0.1939898 | -0.5578372 |
| Uncharacterized protein OS=Schistocephalus solidus OX=70667 GN=SSLN_LOCUS2011 PE=4 SV=1 | 0.54285089 | 0.22968491 | 0.24583586 | -0.7870222 |
| Uncharacterized protein OS=Schistocephalus solidus OX=70667 GN=SSLN_LOCUS9114 PE=4 SV=1 | -0.595353 | -1.3728904 | 0.3993219 | 0.24782257 |
| Cupin domain OS=Schistocephalus solidus OX=70667 GN=SSLN_LOCUS9487 PE=4 SV=1 | -0.4214336 | 0.37401028 | 0.14993893 | -0.5287298 |
| Uncharacterized protein OS=Schistocephalus solidus OX=70667 GN=SSLN_LOCUS18307 PE=3 SV=1 | -0.7869216 | 0.5131663 | 0.15920361 | -0.5169663 |
| 40S ribosomal protein S16 OS=Schistocephalus solidus OX=70667 GN=RS16 PE=3 SV=1 | 0.31299838 | -0.8003206 | 0.79393423 | 0.30077283 |
| Histone H2A OS=Schistocephalus solidus OX=70667 GN=SSLN_LOCUS17647 PE=3 SV=1 | -0.7536063 | -0.9564676 | 0.29160704 | -0.1484811 |
| WD_REPEATS_REGION domain-containing protein OS=Schistocephalus solidus OX=70667 GN=SSLN_LOCUS7520 PE=4 SV=1 | 0.20572858 | -0.8839666 | 0.84824033 | 0.35061948 |
| Uncharacterized protein OS=Schistocephalus solidus OX=70667 GN=SSLN_LOCUS8125 PE=4 SV=1 | -0.0332444 | 0.06855056 | -0.0298743 | -0.7043687 |
| Fibrillar collagen NC1 domain-containing protein OS=Schistocephalus solidus OX=70667 GN=SSLN_LOCUS12411 PE=4 SV=1 | -0.3491311 | 0.17751126 | 0.11574753 | -1.076373 |
| Uncharacterized protein OS=Schistocephalus solidus OX=70667 GN=SSLN_LOCUS17817 PE=3 SV=1 | -0.7086184 | -1.6738744 | 0.20079516 | -0.053568 |
| 40S ribosomal protein S18 OS=Schistocephalus solidus OX=70667 GN=SSLN_LOCUS7468 PE=3 SV=1 | -0.9924845 | -1.4700107 | 0.17937603 | 0.11035864 |
| 40S ribosomal protein S7 OS=Schistocephalus solidus OX=70667 GN=SSLN_LOCUS10505 PE=3 SV=1 | -0.714922 | -1.4714563 | 0.03288253 | 0.19307557 |
| Uncharacterized protein OS=Schistocephalus solidus OX=70667 GN=SSLN_LOCUS5047 PE=4 SV=1 | -0.8022282 | -1.2553537 | 0.23121768 | 0.28943689 |
| Chloride intracellular channel exc-4 OS=Schistocephalus solidus OX=70667 GN=SSLN_LOCUS10784 PE=4 SV=1 | -1.0799894 | 0.03994505 | -0.4242074 | -0.5499483 |
| Uncharacterized protein OS=Schistocephalus solidus OX=70667 GN=SSLN_LOCUS9163 PE=4 SV=1 | 0.17394327 | -0.9706026 | 0.7115485 | 0.26638349 |
| Programmed cell death protein 4 OS=Schistocephalus solidus OX=70667 GN=SSLN_LOCUS8140 PE=3 SV=1 | 0.1291489 | -0.8970389 | 0.85217465 | 0.25458042 |
| Elongation factor Tu; mitochondrial OS=Schistocephalus solidus OX=70667 GN=SSLN_LOCUS17173 PE=3 SV=1 | 0.08741421 | -0.8003173 | 0.79941339 | 0.26250783 |
| Adenine phosphoribosyltransferase OS=Schistocephalus solidus OX=70667 GN=SSLN_LOCUS15665 PE=3 SV=1 | 0.18311445 | 0.8949346 | -0.2011234 | 0.13074472 |
| 40S ribosomal protein S12 OS=Schistocephalus solidus OX=70667 GN=SSLN_LOCUS14540 PE=3 SV=1 | -0.9146834 | -1.8276516 | 0.18450345 | -0.0339304 |
| Ig-like domain-containing protein OS=Schistocephalus solidus OX=70667 GN=SSLN_LOCUS15579 PE=4 SV=1 | 0.54422682 | 0.24392902 | 0.28298027 | -0.6595865 |
| Uncharacterized protein OS=Schistocephalus solidus OX=70667 GN=SSLN_LOCUS6933 PE=4 SV=1 | -0.8676628 | 0.10336112 | 0.00027974 | -0.6734928 |
| Protein transport protein SEC23 OS=Schistocephalus solidus OX=70667 GN=SSLN_LOCUS15867 PE=3 SV=1 | -0.0894988 | -1.0592983 | 0.9710937 | 0.39335941 |
| Uncharacterized protein OS=Schistocephalus solidus OX=70667 GN=SSLN_LOCUS1400 PE=4 SV=1 | 0.70585895 | 0.12710785 | 0.28399832 | -0.6065008 |
| Glutamine synthetase OS=Schistocephalus solidus OX=70667 GN=SSLN_LOCUS2464 PE=3 SV=1 | 0.05702007 | -0.9114233 | 0.89025694 | 0.14834351 |
| EF-hand domain-containing protein OS=Schistocephalus solidus OX=70667 GN=SSLN_LOCUS2894 PE=4 SV=1 | -0.4109782 | 0.73338446 | 0.35715219 | -0.5417914 |
| Histone H2A OS=Schistocephalus solidus OX=70667 GN=SSLN_LOCUS17074 PE=3 SV=1 | -0.7810675 | -1.2293995 | 0.44123586 | 0.21581501 |
| Ribosomal protein L15 OS=Schistocephalus solidus OX=70667 GN=SSLN_LOCUS18365 PE=3 SV=1 | 0.16305834 | -0.9676877 | 0.73278481 | 0.25061293 |
| KH domain-containing protein OS=Schistocephalus solidus OX=70667 GN=SSLN_LOCUS16010 PE=4 SV=1 | 0.44076064 | -0.7525272 | 0.77950986 | 0.50058387 |
| L-threonine 3-dehydrogenase OS=Schistocephalus solidus OX=70667 GN=SSLN_LOCUS7096 PE=4 SV=1 | 1.45610324 | -0.2074911 | 0.10750594 | 0.59772786 |
| Purine nucleoside phosphorylase OS=Schistocephalus solidus OX=70667 GN=SSLN_LOCUS14361 PE=3 SV=1 | -1.0667552 | -0.0311752 | -0.5164492 | 0.55289858 |
| Histone H2A OS=Schistocephalus solidus OX=70667 GN=H2AX PE=3 SV=1 | -0.9489631 | -1.0238125 | 0.32147666 | -0.1610709 |
| Uncharacterized protein OS=Schistocephalus solidus OX=70667 GN=SSLN_LOCUS9256 PE=4 SV=1 | 1.23147194 | -0.1152327 | 0.23494389 | -0.5558282 |
| RRM domain-containing protein OS=Schistocephalus solidus OX=70667 GN=SSLN_LOCUS11874 PE=4 SV=1 | -0.9395177 | -1.8312002 | 0.20641638 | -0.0473206 |
| ARMET_N domain-containing protein OS=Schistocephalus solidus OX=70667 GN=SSLN_LOCUS4913 PE=4 SV=1 | -0.2801187 | -1.1210608 | 1.00655064 | 0.31970533 |
| ATP synthase subunit O; mitochondrial OS=Schistocephalus solidus OX=70667 GN=SSLN_LOCUS11656 PE=3 SV=1 | 0.31516383 | -0.7867357 | 0.78447068 | 0.42277709 |
| SERPIN domain-containing protein OS=Schistocephalus solidus OX=70667 GN=SSLN_LOCUS18505 PE=3 SV=1 | -0.0618704 | 0.9324213 | 0.06441879 | 0.02797257 |
| Uncharacterized protein OS=Schistocephalus solidus OX=70667 GN=SSLN_LOCUS3219 PE=4 SV=1 | 0.00518351 | 0.36629681 | 0.35428611 | -0.6728594 |
| 60S ribosomal protein L36a OS=Schistocephalus solidus OX=70667 GN=RL36A PE=3 SV=1 | 0.45838656 | -0.7216297 | 0.78920736 | 0.50910131 |
| Uncharacterized protein OS=Schistocephalus solidus OX=70667 GN=SSLN_LOCUS18466 PE=4 SV=1 | 1.43959015 | -0.3164047 | 0.26867392 | 0.57944623 |
| Uncharacterized protein OS=Schistocephalus solidus OX=70667 GN=SSLN_LOCUS18935 PE=4 SV=1 | -0.0275884 | 0.6621181 | 0.29909381 | -0.7370519 |
| Splicing factor U2AF subunit OS=Schistocephalus solidus OX=70667 GN=U2AF2 PE=3 SV=1 | 1.88515272 | -0.245591 | -0.5485398 | 0.15224256 |
| Alpha-carbonic anhydrase domain-containing protein OS=Schistocephalus solidus OX=70667 GN=SSLN_LOCUS8517 PE=3 SV=1 | -0.5285252 | -0.0174787 | -0.1432364 | -1.1272247 |
| Mitotic checkpoint protein BUB3 OS=Schistocephalus solidus OX=70667 GN=SSLN_LOCUS135 PE=4 SV=1 | 0.38381893 | -0.7163951 | 0.82636402 | 0.24519812 |
| Uncharacterized protein OS=Schistocephalus solidus OX=70667 GN=SSLN_LOCUS4559 PE=4 SV=1 | -0.8394896 | 0.39516761 | 0.09774015 | -0.9536477 |
| Pribosyltran domain-containing protein OS=Schistocephalus solidus OX=70667 GN=SSLN_LOCUS17289 PE=4 SV=1 | -1.5509387 | -0.0898879 | -0.9495318 | 0.17793885 |
| Transmembrane emp24 domain-containing protein 2 OS=Schistocephalus solidus OX=70667 GN=TMED2 PE=3 SV=1 | -0.3621784 | -1.1882781 | 1.1378428 | 0.42675004 |
| JAB_MPN domain-containing protein OS=Schistocephalus solidus OX=70667 GN=SSLN_LOCUS11330 PE=4 SV=1 | 1.9023577 | -0.2777077 | -0.5573765 | 0.18410665 |
| Uncharacterized protein OS=Schistocephalus solidus OX=70667 GN=SSLN_LOCUS15939 PE=3 SV=1 | 0.226886 | 0.90064201 | -0.2078952 | 0.21217983 |
| Eukaryotic translation initiation factor 3 subunit B OS=Schistocephalus solidus OX=70667 GN=SSLN_LOCUS15481 PE=3 SV=1 | 0.05932311 | -0.9320106 | 1.03347884 | 0.60594452 |
| Uncharacterized protein OS=Schistocephalus solidus OX=70667 GN=SSLN_LOCUS5487 PE=4 SV=1 | -0.1820568 | 0.31460527 | 0.34912585 | 0.54303715 |
| Uncharacterized protein OS=Schistocephalus solidus OX=70667 GN=SSLN_LOCUS9932 PE=4 SV=1 | -0.1174768 | 0.8971885 | 0.21048657 | 0.07796808 |
| Uncharacterized protein OS=Schistocephalus solidus OX=70667 GN=SSLN_LOCUS19196 PE=4 SV=1 | -0.1940744 | 0.66710304 | 0.27648729 | -0.7698339 |
| Uncharacterized protein OS=Schistocephalus solidus OX=70667 GN=SSLN_LOCUS16625 PE=4 SV=1 | -0.0600406 | 1.05143898 | -0.185672 | 0.16626114 |
| Sm domain-containing protein OS=Schistocephalus solidus OX=70667 GN=SSLN_LOCUS12897 PE=4 SV=1 | 1.48022903 | -0.3064347 | 0.20521077 | 0.55391605 |
| Uncharacterized protein OS=Schistocephalus solidus OX=70667 GN=SSLN_LOCUS14721 PE=4 SV=1 | -0.0432696 | 0.47799375 | 0.45303137 | -1.0136621 |
| Uncharacterized protein OS=Schistocephalus solidus OX=70667 GN=SSLN_LOCUS2366 PE=4 SV=1 | -0.2939213 | 0.47477907 | 0.50383582 | -0.9441503 |
| Adenylyl-sulfate kinase OS=Schistocephalus solidus OX=70667 GN=SSLN_LOCUS14484 PE=3 SV=1 | 0.82182325 | 0.67076671 | -0.2902451 | 0.62774137 |
| Uncharacterized protein OS=Schistocephalus solidus OX=70667 GN=SSLN_LOCUS7964 PE=4 SV=1 | 1.64309194 | 0.39174753 | -0.8131891 | 0.26682246 |
| Uncharacterized protein OS=Schistocephalus solidus OX=70667 GN=SSLN_LOCUS4572 PE=4 SV=1 | -0.1712464 | 1.12553633 | -0.2525896 | 0.20729366 |
| HABP4_PAI-RBP1 domain-containing protein OS=Schistocephalus solidus OX=70667 GN=SSLN_LOCUS12473 PE=4 SV=1 | -1.4418171 | -2.133186 | -0.1313972 | -0.0247426 |
| Uncharacterized protein OS=Schistocephalus solidus OX=70667 GN=SSLN_LOCUS10621 PE=4 SV=1 | 1.43549231 | -0.3051853 | 0.29235785 | 0.55819343 |
| Aldedh domain-containing protein (Fragment) OS=Schistocephalus solidus OX=70667 GN=SSLN_LOCUS18129 PE=4 SV=1 | 1.4429213 | -0.2621035 | 0.25881063 | 0.5859993 |
| Ste24 endopeptidase OS=Schistocephalus solidus OX=70667 GN=SSLN_LOCUS15092 PE=4 SV=1 | -0.4486668 | -1.2904183 | 0.97681143 | 0.14907192 |
| Uncharacterized protein OS=Schistocephalus solidus OX=70667 GN=SSLN_LOCUS5672 PE=4 SV=1 | 1.24304309 | -0.1146716 | 0.20091735 | -0.594351 |
| PlsC domain-containing protein OS=Schistocephalus solidus OX=70667 GN=SSLN_LOCUS8425 PE=3 SV=1 | 1.49967919 | -0.2700678 | 0.16831889 | 0.54072469 |
| Uncharacterized protein OS=Schistocephalus solidus OX=70667 GN=SSLN_LOCUS13395 PE=3 SV=1 | 1.43023928 | -0.2076217 | 0.18769562 | 0.66813342 |
| Uncharacterized protein OS=Schistocephalus solidus OX=70667 GN=SSLN_LOCUS13073 PE=4 SV=1 | 1.37701213 | -0.1554514 | 0.08166349 | -0.5247782 |
| ATP citrate synthase OS=Schistocephalus solidus OX=70667 GN=SSLN_LOCUS6335 PE=4 SV=1 | 1.72872017 | 0.23532606 | -0.8106995 | 0.34574231 |
| Uncharacterized protein OS=Schistocephalus solidus OX=70667 GN=SSLN_LOCUS6165 PE=4 SV=1 | 1.57391426 | 0.50005929 | -0.8067423 | 0.20927778 |
| SPRY domain-containing protein OS=Schistocephalus solidus OX=70667 GN=SSLN_LOCUS6259 PE=4 SV=1 | 1.42541782 | -0.1190107 | -0.0447346 | -0.5405672 |

| Glyoxalase domain-containing protein 4 OS=Schistocephalus solidus OX=70667 GN=SSLN_LOCUS13951 PE=3 SV=1 | 0.79958675 | 0.92885185 | -0.6046435 | 0.34510756 |
| --- | --- | --- | --- | --- |
| Uncharacterized protein OS=Schistocephalus solidus OX=70667 GN=SSLN_LOCUS970 PE=4 SV=1 | 1.87787819 | -0.2556177 | -0.5347452 | 0.09510145 |
| TPR_REGION domain-containing protein OS=Schistocephalus solidus OX=70667 GN=SSLN_LOCUS4410 PE=4 SV=1 | 1.69858104 | -0.3121508 | -0.1669346 | 0.51623402 |
| MYND-type domain-containing protein OS=Schistocephalus solidus OX=70667 GN=SSLN_LOCUS4395 PE=4 SV=1 | 1.90542923 | -0.2722671 | -0.5548445 | 0.16172422 |
| Aldedh domain-containing protein OS=Schistocephalus solidus OX=70667 GN=SSLN_LOCUS17637 PE=4 SV=1 | 1.63384372 | 0.35787735 | -0.7497584 | 0.26238043 |
| Uncharacterized protein OS=Schistocephalus solidus OX=70667 GN=SSLN_LOCUS3031 PE=3 SV=1 | 1.21677825 | -0.0929007 | 0.21571039 | -0.6523461 |
| Uncharacterized protein OS=Schistocephalus solidus OX=70667 GN=SSLN_LOCUS8454 PE=3 SV=1 | 1.41206896 | -0.1285631 | -0.0231873 | -0.5831138 |
| tRNA-synt_1c domain-containing protein OS=Schistocephalus solidus OX=70667 GN=SSLN_LOCUS17070 PE=3 SV=1 | 1.88017031 | -0.248703 | -0.5380897 | 0.11392449 |
| Uncharacterized protein OS=Schistocephalus solidus OX=70667 GN=SSLN_LOCUS10025 PE=4 SV=1 | -0.053064 | 0.98775212 | -0.1552659 | 0.13376438 |
| 60S ribosomal protein L31 OS=Schistocephalus solidus OX=70667 GN=SSLN_LOCUS14236 PE=3 SV=1 | 1.90237524 | -0.2865457 | -0.5489475 | 0.13833661 |
| Uncharacterized protein OS=Schistocephalus solidus OX=70667 GN=SSLN_LOCUS8980 PE=4 SV=1 | 1.88247248 | -0.2781465 | -0.5072579 | 0.13455659 |
| Uncharacterized protein OS=Schistocephalus solidus OX=70667 GN=SSLN_LOCUS1729 PE=4 SV=1 | -0.311068 | 0.56906388 | 0.43888053 | -1.2556371 |
| Uncharacterized protein OS=Schistocephalus solidus OX=70667 GN=SSLN_LOCUS16490 PE=4 SV=1 | -0.212936 | 0.53417968 | 0.39238444 | -1.165468 |
| Uncharacterized protein OS=Schistocephalus solidus OX=70667 GN=SSLN_LOCUS4232 PE=4 SV=1 | -0.1957388 | 1.05602912 | 0.06672541 | 0.3158905 |
| F-actin-capping protein subunit alpha OS=Schistocephalus solidus OX=70667 GN=SSLN_LOCUS52 PE=3 SV=1 | 0.14740982 | 1.02671234 | -0.2646473 | 0.17420964 |
| Uncharacterized protein OS=Schistocephalus solidus OX=70667 GN=SSLN_LOCUS12448 PE=4 SV=1 | 1.8755014 | -0.2419031 | -0.5248893 | 0.1167642 |
| DZF domain-containing protein OS=Schistocephalus solidus OX=70667 GN=SSLN_LOCUS6401 PE=4 SV=1 | 1.89890164 | -0.3073923 | -0.5138347 | 0.18641679 |
| Peptidase A1 domain-containing protein OS=Schistocephalus solidus OX=70667 GN=SSLN_LOCUS3359 PE=3 SV=1 | 1.47819531 | 0.83235832 | -1.0233186 | 0.49255657 |
| Uncharacterized protein OS=Schistocephalus solidus OX=70667 GN=SSLN_LOCUS2561 PE=3 SV=1 | 1.87785851 | -0.2484882 | -0.5292595 | 0.11371991 |
| Fibrillar collagen NC1 domain-containing protein OS=Schistocephalus solidus OX=70667 GN=SSLN_LOCUS15111 PE=4 SV=1 | -0.2214911 | 0.53250566 | 0.43331852 | -1.1401566 |
| CS domain-containing protein OS=Schistocephalus solidus OX=70667 GN=SSLN_LOCUS11757 PE=4 SV=1 | 1.88151422 | -0.2572366 | -0.544312 | 0.10064232 |
| Uncharacterized protein OS=Schistocephalus solidus OX=70667 GN=SSLN_LOCUS7620 PE=4 SV=1 | 0.7854076 | 0.92551263 | -0.606093 | 0.33970466 |
| Cystatin domain-containing protein OS=Schistocephalus solidus OX=70667 GN=SSLN_LOCUS18362 PE=4 SV=1 | -0.6849987 | 0.90352446 | 0.40376102 | -0.6274428 |
| RRM domain-containing protein OS=Schistocephalus solidus OX=70667 GN=SSLN_LOCUS624 PE=4 SV=1 | 1.88840214 | -0.2631767 | -0.531298 | 0.1447895 |
| ATP synthase-coupling factor 6; mitochondrial OS=Schistocephalus solidus OX=70667 GN=SSLN_LOCUS1516 PE=3 SV=1 | 1.75718232 | 0.12154283 | -0.7361525 | 0.30633898 |
| Uncharacterized protein OS=Schistocephalus solidus OX=70667 GN=SSLN_LOCUS4126 PE=3 SV=1 | 1.89910022 | -0.2720652 | -0.5437128 | 0.1602991 |
| Uncharacterized protein OS=Schistocephalus solidus OX=70667 GN=SSLN_LOCUS4976 PE=4 SV=1 | 1.88711661 | -0.2513435 | -0.55165 | 0.11893835 |
| Coatomer subunit epsilon OS=Schistocephalus solidus OX=70667 GN=SSLN_LOCUS9062 PE=3 SV=1 | 1.87734717 | -0.242354 | -0.5445709 | 0.10376687 |
| Uncharacterized protein OS=Schistocephalus solidus OX=70667 GN=SSLN_LOCUS8649 PE=4 SV=1 | 1.92549702 | -0.2886957 | -0.5945388 | 0.21347388 |
| Uncharacterized protein OS=Schistocephalus solidus OX=70667 GN=SSLN_LOCUS13285 PE=4 SV=1 | 0.19733472 | 0.88110769 | -0.3927803 | 0.17795435 |
| Tr-type G domain-containing protein OS=Schistocephalus solidus OX=70667 GN=SSLN_LOCUS19932 PE=4 SV=1 | 1.7000887 | 0.27926233 | -0.803644 | 0.2825015 |
| Mitochondrial import receptor subunit TOM22 homolog OS=Schistocephalus solidus OX=70667 GN=SSLN_LOCUS15875 PE=3 SV=1 | 1.44762367 | -0.3420325 | 0.24521255 | 0.74955855 |
| Phosducin domain-containing protein OS=Schistocephalus solidus OX=70667 GN=SSLN_LOCUS7877 PE=3 SV=1 | 0.87220579 | 0.88833939 | -0.6772387 | 0.44579336 |
| Uncharacterized protein OS=Schistocephalus solidus OX=70667 GN=SSLN_LOCUS10842 PE=4 SV=1 | 1.89904169 | -0.2779537 | -0.5636994 | 0.1295515 |
| Uncharacterized protein OS=Schistocephalus solidus OX=70667 GN=SSLN_LOCUS19531 PE=4 SV=1 | 1.88798021 | -0.2629549 | -0.5491895 | 0.14350126 |
| Dolichol-phosphate mannosyltransferase subunit 1 OS=Schistocephalus solidus OX=70667 GN=SSLN_LOCUS3702 PE=3 SV=1 | 1.54542914 | -0.2868656 | 0.11957749 | 0.53627093 |
| Uncharacterized protein OS=Schistocephalus solidus OX=70667 GN=SSLN_LOCUS7085 PE=4 SV=1 | -0.5781188 | 0.88046622 | 0.33841067 | -0.7921278 |
| GRASP55_65 domain-containing protein OS=Schistocephalus solidus OX=70667 GN=SSLN_LOCUS7795 PE=3 SV=1 | 1.91518011 | -0.2936019 | -0.5695399 | 0.20421764 |
| Coatomer subunit delta OS=Schistocephalus solidus OX=70667 GN=SSLN_LOCUS11855 PE=3 SV=1 | 1.91215167 | -0.258263 | -0.5748499 | 0.15371326 |
| Nop domain-containing protein OS=Schistocephalus solidus OX=70667 GN=SSLN_LOCUS11486 PE=3 SV=1 | 1.90967503 | -0.2806856 | -0.5718166 | 0.14338877 |
| Uncharacterized protein OS=Schistocephalus solidus OX=70667 GN=SSLN_LOCUS9677 PE=3 SV=1 | 1.91518011 | -0.2936019 | -0.5695399 | 0.20421764 |
| Uncharacterized protein OS=Schistocephalus solidus OX=70667 GN=SSLN_LOCUS42 PE=4 SV=1 | 0.8113067 | 1.00252531 | -0.7099119 | 0.43774354 |
| Uncharacterized protein OS=Schistocephalus solidus OX=70667 GN=SSLN_LOCUS342 PE=4 SV=1 | 1.91218324 | -0.2745575 | -0.5753462 | 0.10867796 |
| Uncharacterized protein OS=Schistocephalus solidus OX=70667 GN=SSLN_LOCUS222 PE=4 SV=1 | 1.90177485 | -0.2834406 | -0.5630094 | 0.15618444 |
| Myosin motor domain-containing protein OS=Schistocephalus solidus OX=70667 GN=SSLN_LOCUS12218 PE=3 SV=1 | 1.55393266 | 0.66141121 | -0.9778222 | 0.45739772 |
| HELP domain-containing protein OS=Schistocephalus solidus OX=70667 GN=SSLN_LOCUS2156 PE=3 SV=1 | 1.56051364 | -0.0100941 | -0.2328181 | 0.52767603 |
| Splicing factor 3A subunit 1 OS=Schistocephalus solidus OX=70667 GN=SSLN_LOCUS4535 PE=4 SV=1 | 1.39329979 | -0.3178084 | 0.30386555 | 0.72808032 |
| Phosphotransferase OS=Schistocephalus solidus OX=70667 GN=SSLN_LOCUS6040 PE=3 SV=1 | 1.41013014 | -0.3430105 | 0.3067705 | 0.80400937 |
| Uncharacterized protein OS=Schistocephalus solidus OX=70667 GN=SSLN_LOCUS8719 PE=4 SV=1 | -0.1885608 | 1.0769167 | -0.1512489 | 0.19275515 |
| Queuosine salvage protein OS=Schistocephalus solidus OX=70667 GN=SSLN_LOCUS8908 PE=3 SV=1 | 1.73114177 | 0.22306126 | -0.8016986 | 0.34969297 |
| Uncharacterized protein OS=Schistocephalus solidus OX=70667 GN=SSLN_LOCUS11736 PE=3 SV=1 | 1.90537504 | -0.2529567 | -0.5675055 | 0.12110337 |
| Uncharacterized protein OS=Schistocephalus solidus OX=70667 GN=SSLN_LOCUS14207 PE=4 SV=1 | 1.87332678 | -0.2829737 | -0.5077324 | 0.07839395 |
| Uncharacterized protein OS=Schistocephalus solidus OX=70667 GN=SSLN_LOCUS15791 PE=4 SV=1 | 1.6919396 | 0.27351371 | -0.7838967 | 0.31867479 |
| J domain-containing protein OS=Schistocephalus solidus OX=70667 GN=SSLN_LOCUS18257 PE=4 SV=1 | 1.43046395 | -0.3540932 | 0.26463448 | 0.70900051 |
| Importin subunit beta-1 OS=Schistocephalus solidus OX=70667 GN=SSLN_LOCUS5906 PE=4 SV=1 | 1.91696324 | -0.2900543 | -0.595551 | 0.24341983 |
| CRAL_TRIO_N domain-containing protein OS=Schistocephalus solidus OX=70667 GN=SSLN_LOCUS13352 PE=4 SV=1 | 0.56572936 | 1.12817697 | -0.5595386 | 0.48888852 |
| PP1-binding domain-containing protein OS=Schistocephalus solidus OX=70667 GN=SSLN_LOCUS15190 PE=4 SV=1 | 1.89698807 | -0.285004 | -0.5281632 | 0.08943936 |
| Ectonucleotide pyrophosphatase/phosphodiesterase family member 5 OS=Schistocephalus solidus OX=70667 GN=SSLN_LOCUS8008 PE=4 SV=1 | 1.59581752 | 0.5668727 | -0.9526615 | 0.43795391 |
| V-SNARE coiled-coil homology domain-containing protein OS=Schistocephalus solidus OX=70667 GN=SSLN_LOCUS6267 PE=4 SV=1 | 1.47819531 | 0.83235832 | -1.0233186 | 0.49255657 |
| Uncharacterized protein OS=Schistocephalus solidus OX=70667 GN=SSLN_LOCUS9969 PE=4 SV=1 | 1.91921872 | -0.2932679 | -0.578173 | 0.21480309 |
| EGF-like domain-containing protein OS=Schistocephalus solidus OX=70667 GN=SSLN_LOCUS14554 PE=4 SV=1 | 1.91921872 | -0.2932679 | -0.578173 | 0.21480309 |
| Nucleolar protein 10 OS=Schistocephalus solidus OX=70667 GN=SSLN_LOCUS12853 PE=3 SV=1 | 1.90278987 | -0.2481192 | -0.5778329 | 0.13338046 |
| N-acetyltransferase domain-containing protein OS=Schistocephalus solidus OX=70667 GN=SSLN_LOCUS17678 PE=4 SV=1 | 1.91921872 | -0.2932679 | -0.578173 | 0.21480309 |
| H/ACA ribonucleoprotein complex subunit OS=Schistocephalus solidus OX=70667 GN=SSLN_LOCUS5277 PE=3 SV=1 | 1.61142635 | -0.3215967 | -0.0077486 | 0.64029545 |
| Pyridoxal phosphate homeostasis protein OS=Schistocephalus solidus OX=70667 GN=SSLN_LOCUS10286 PE=3 SV=1 | 1.7448909 | 0.13078119 | -0.7427817 | 0.25392569 |
| Dolichyl-diphosphooligosaccharide--protein glycotransferase OS=Schistocephalus solidus OX=70667 GN=SSLN_LOCUS12804 PE=3 SV=1 | 1.90731232 | -0.2733193 | -0.5733513 | 0.10700463 |
| Uncharacterized protein OS=Schistocephalus solidus OX=70667 GN=SSLN_LOCUS18166 PE=3 SV=1 | 1.61746097 | -0.3166381 | -0.0115796 | 0.64172483 |
| CTP:phosphoethanolamine cytidylyltransferase OS=Schistocephalus solidus OX=70667 GN=SSLN_LOCUS6435 PE=3 SV=1 | 0.04187822 | 1.07984478 | -0.4094707 | 0.29154396 |
| FERM domain-containing protein OS=Schistocephalus solidus OX=70667 GN=SSLN_LOCUS14221 PE=4 SV=1 | 0.05776958 | 1.0901671 | -0.4227028 | 0.29755287 |
| ATP-grasp_2 domain-containing protein OS=Schistocephalus solidus OX=70667 GN=SSLN_LOCUS17550 PE=4 SV=1 | 1.36771172 | -0.0218579 | 0.07227861 | 0.68228155 |
| Uncharacterized protein OS=Schistocephalus solidus OX=70667 GN=SSLN_LOCUS14042 PE=4 SV=1 | 0.79713249 | 0.92657883 | -0.5346094 | 0.4268225 |
| Uncharacterized protein OS=Schistocephalus solidus OX=70667 GN=SSLN_LOCUS272 PE=4 SV=1 | 0.57145839 | 1.01829548 | -0.4594246 | 0.43107834 |
| Peptidase_M28 domain-containing protein OS=Schistocephalus solidus OX=70667 GN=SSLN_LOCUS10325 PE=4 SV=1 | 1.74909749 | 0.16538686 | -0.7761222 | 0.36262359 |
| Clu domain-containing protein OS=Schistocephalus solidus OX=70667 GN=SSLN_LOCUS4674 PE=4 SV=1 | 1.94515093 | -0.2866315 | -0.6247472 | 0.23267469 |
| 26S proteasome non-ATPase regulatory subunit 5 OS=Schistocephalus solidus OX=70667 GN=SSLN_LOCUS18236 PE=3 SV=1 | 1.89845886 | -0.2491165 | -0.5690737 | 0.16468055 |
| Signal recognition particle 54 kDa protein OS=Schistocephalus solidus OX=70667 GN=SRP54 PE=3 SV=1 | 1.94472941 | -0.2889967 | -0.6127643 | 0.2182637 |
| Uncharacterized protein OS=Schistocephalus solidus OX=70667 GN=SSLN_LOCUS10716 PE=4 SV=1 | 1.63434784 | 0.41667082 | -0.8605297 | 0.38899055 |
| Uncharacterized protein OS=Schistocephalus solidus OX=70667 GN=SSLN_LOCUS5974 PE=4 SV=1 | -0.000808 | 1.18779958 | -0.4278353 | 0.33306735 |
| C2H2-type domain-containing protein OS=Schistocephalus solidus OX=70667 GN=SSLN_LOCUS12336 PE=4 SV=1 | 1.90969299 | -0.3099664 | -0.5205869 | 0.19533086 |
| Septin-type G domain-containing protein OS=Schistocephalus solidus OX=70667 GN=SSLN_LOCUS19286 PE=4 SV=1 | 1.94101756 | -0.2951055 | -0.6010044 | 0.23497194 |
| LSDAT_euk domain-containing protein OS=Schistocephalus solidus OX=70667 GN=SSLN_LOCUS16364 PE=4 SV=1 | -0.0079216 | 1.01732548 | -0.3810184 | 0.22461599 |
| Uncharacterized protein OS=Schistocephalus solidus OX=70667 GN=SSLN_LOCUS16829 PE=4 SV=1 | 0.51048408 | 1.26406524 | -0.6845284 | 0.51274681 |
| Uncharacterized protein OS=Schistocephalus solidus OX=70667 GN=SSLN_LOCUS971 PE=4 SV=1 | 1.94393577 | -0.2895166 | -0.6152583 | 0.22950738 |
| Uncharacterized protein OS=Schistocephalus solidus OX=70667 GN=SSLN_LOCUS4602 PE=4 SV=1 | 1.73089171 | 0.24666971 | -0.8249833 | 0.3277324 |
| DZF domain-containing protein OS=Schistocephalus solidus OX=70667 GN=SSLN_LOCUS6169 PE=4 SV=1 | 1.94310511 | -0.2930064 | -0.5991043 | 0.21253471 |
| Uncharacterized protein OS=Schistocephalus solidus OX=70667 GN=SSLN_LOCUS6469 PE=4 SV=1 | 0.71761189 | 1.00731879 | -0.520939 | 0.43760404 |
| Geranylgeranyl transferase type-2 subunit alpha OS=Schistocephalus solidus OX=70667 GN=SSLN_LOCUS8418 PE=3 SV=1 | 1.73118878 | 0.21795124 | -0.8053536 | 0.37185024 |
| Uncharacterized protein OS=Schistocephalus solidus OX=70667 GN=SSLN_LOCUS8907 PE=3 SV=1 | 1.94310511 | -0.2930064 | -0.5991043 | 0.21253471 |
| ATP citrate synthase OS=Schistocephalus solidus OX=70667 GN=SSLN_LOCUS9514 PE=4 SV=1 | 0.65286977 | 1.13347436 | -0.6991632 | 0.49335616 |
| Uncharacterized protein OS=Schistocephalus solidus OX=70667 GN=SSLN_LOCUS10624 PE=4 SV=1 | 0.66170041 | 1.11826641 | -0.6745935 | 0.44939659 |
| Complex1_LYR_dom domain-containing protein OS=Schistocephalus solidus OX=70667 GN=SSLN_LOCUS10996 PE=3 SV=1 | 1.55471442 | -0.3211108 | 0.09398739 | 0.67129709 |
| TPR_REGION domain-containing protein OS=Schistocephalus solidus OX=70667 GN=SSLN_LOCUS12812 PE=4 SV=1 | 1.56744655 | -0.3196281 | 0.06753819 | 0.72896119 |
| Thioredoxin domain-containing protein OS=Schistocephalus solidus OX=70667 GN=SSLN_LOCUS15840 PE=4 SV=1 | 1.94393577 | -0.2895166 | -0.6152583 | 0.22950738 |
| Uncharacterized protein OS=Schistocephalus solidus OX=70667 GN=SSLN_LOCUS16249 PE=4 SV=1 | -1.2154898 | 0.28144464 | 0.17790743 | -1.4592035 |
| Uncharacterized protein OS=Schistocephalus solidus OX=70667 GN=SSLN_LOCUS16878 PE=4 SV=1 | 1.94393577 | -0.2895166 | -0.6152583 | 0.22950738 |
| PI31_Prot_C domain-containing protein OS=Schistocephalus solidus OX=70667 GN=SSLN_LOCUS19472 PE=3 SV=1 | 1.94310511 | -0.2930064 | -0.5991043 | 0.21253471 |
| ER membrane protein complex subunit 1 OS=Schistocephalus solidus OX=70667 GN=SSLN_LOCUS9557 PE=3 SV=1 | 1.8958568 | -0.308714 | -0.5093425 | 0.19013384 |
| protein-synthesizing GTPase OS=Caenorhabditis elegans OX=6239 GN=eif-2gamma PE=1 SV=1 | -0.6969395 | 0.7788194 | 0.47681374 | 0.53202106 |
| Importin N-terminal domain-containing protein OS=Schistocephalus solidus OX=70667 GN=SSLN_LOCUS12808 PE=3 SV=1 | 1.95527997 | -0.2692139 | -0.6616 | 0.19437424 |
| Uncharacterized protein OS=Schistocephalus solidus OX=70667 GN=SSLN_LOCUS3175 PE=3 SV=1 | 0.59504953 | 1.22633964 | -0.7141343 | 0.51098245 |
| SH3 domain-containing protein OS=Schistocephalus solidus OX=70667 GN=SSLN_LOCUS10140 PE=4 SV=1 | 0.37577772 | 1.32415927 | -0.6373685 | 0.51555732 |
| Arrestin_N domain-containing protein OS=Schistocephalus solidus OX=70667 GN=SSLN_LOCUS14326 PE=4 SV=1 | -0.3997017 | 0.92139241 | 0.49641315 | 0.37236562 |
| Beta-N-acetylhexosaminidase OS=Schistocephalus solidus OX=70667 GN=SSLN_LOCUS8403 PE=3 SV=1 | 1.95919047 | -0.253298 | -0.7343788 | 0.26926865 |
| Uncharacterized protein OS=Schistocephalus solidus OX=70667 GN=SSLN_LOCUS8724 PE=4 SV=1 | 1.47819531 | 0.83235832 | -1.0233186 | 0.49255657 |
| SH3 domain-containing protein OS=Schistocephalus solidus OX=70667 GN=SSLN_LOCUS12984 PE=4 SV=1 | 1.47819531 | 0.83235832 | -1.0233186 | 0.49255657 |
| Uncharacterized protein OS=Schistocephalus solidus OX=70667 GN=SSLN_LOCUS730 PE=4 SV=1 | 0.40569998 | 1.31081061 | -0.6478441 | 0.51493302 |
| Density-regulated protein OS=Schistocephalus solidus OX=70667 GN=SSLN_LOCUS2350 PE=4 SV=1 | 1.59719231 | 0.55155364 | -0.9147427 | 0.39944175 |
| Uncharacterized protein OS=Schistocephalus solidus OX=70667 GN=SSLN_LOCUS5759 PE=3 SV=1 | 1.95462346 | -0.2718859 | -0.6493817 | 0.18180076 |
| SH3 domain-containing protein OS=Schistocephalus solidus OX=70667 GN=SSLN_LOCUS6546 PE=4 SV=1 | 0.54389542 | 1.24916006 | -0.6962255 | 0.51204972 |
| Protein kinase domain-containing protein OS=Schistocephalus solidus OX=70667 GN=SSLN_LOCUS9638 PE=3 SV=1 | 0.54389542 | 1.24916006 | -0.6962255 | 0.51204972 |
| Glutamate-5-semialdehyde dehydrogenase OS=Schistocephalus solidus OX=70667 GN=SSLN_LOCUS9836 PE=3 SV=1 | 1.94095271 | -0.2914708 | -0.6246327 | 0.27176926 |
| Uncharacterized protein OS=Schistocephalus solidus OX=70667 GN=SSLN_LOCUS12841 PE=4 SV=1 | 1.54108417 | 0.59619054 | -0.8288545 | 0.24196865 |
| Arginyltransferase OS=Schistocephalus solidus OX=70667 GN=SSLN_LOCUS16715 PE=3 SV=1 | 1.59581911 | 0.5668691 | -0.9526605 | 0.43795316 |
| NEDD8 OS=Schistocephalus solidus OX=70667 GN=SSLN_LOCUS18831 PE=3 SV=1 | 1.54108417 | 0.59619054 | -0.8288545 | 0.24196865 |
| Methionyl-tRNA synthetase OS=Schistocephalus solidus OX=70667 GN=SSLN_LOCUS19743 PE=3 SV=1 | 1.88504958 | -0.2349985 | -0.6237635 | 0.04636492 |
| Signal peptidase complex catalytic subunit SEC11 OS=Schistocephalus solidus OX=70667 GN=SSLN_LOCUS7161 PE=3 SV=1 | 1.87359597 | -0.2119758 | -0.5347623 | -0.0020925 |
| Uncharacterized protein OS=Schistocephalus solidus OX=70667 GN=SSLN_LOCUS6353 PE=3 SV=1 | 1.95462346 | -0.2718859 | -0.6493817 | 0.18180076 |
| Lactamase_B domain-containing protein OS=Schistocephalus solidus OX=70667 GN=SSLN_LOCUS9380 PE=4 SV=1 | 1.54108417 | 0.59619054 | -0.8288545 | 0.24196865 |
| Uncharacterized protein OS=Schistocephalus solidus OX=70667 GN=SSLN_LOCUS7062 PE=3 SV=1 | -0.0947411 | 1.02813133 | -0.343495 | 0.2089628 |
| Eukaryotic translation initiation factor 3 subunit K OS=Schistocephalus solidus OX=70667 GN=SSLN_LOCUS16656 PE=3 SV=1 | 1.95594064 | -0.266525 | -0.6738958 | 0.20702743 |
| S1 motif domain-containing protein OS=Schistocephalus solidus OX=70667 GN=SSLN_LOCUS12224 PE=4 SV=1 | 1.88909096 | -0.3224859 | -0.4856738 | 0.23136524 |
| Purple acid phosphatase OS=Schistocephalus solidus OX=70667 GN=SSLN_LOCUS6511 PE=3 SV=1 | 1.90637909 | -0.2772356 | -0.6099465 | 0.22343949 |
| Ubiquitin domain-containing protein UBFD1 OS=Schistocephalus solidus OX=70667 GN=UBFD1 PE=4 SV=1 | 1.47819531 | 0.83235832 | -1.0233186 | 0.49255657 |
| Asparaginase OS=Schistocephalus solidus OX=70667 GN=SSLN_LOCUS2489 PE=4 SV=1 | 0.21952425 | 1.39386571 | -0.582665 | 0.51881737 |
| DUF5733 domain-containing protein OS=Schistocephalus solidus OX=70667 GN=SSLN_LOCUS5334 PE=4 SV=1 | 1.47819531 | 0.83235832 | -1.0233186 | 0.49255657 |
| Uncharacterized protein OS=Schistocephalus solidus OX=70667 GN=SSLN_LOCUS5579 PE=4 SV=1 | 1.47819531 | 0.83235832 | -1.0233186 | 0.49255657 |
| Uncharacterized protein OS=Schistocephalus solidus OX=70667 GN=SSLN_LOCUS10764 PE=4 SV=1 | 1.47819531 | 0.83235832 | -1.0233186 | 0.49255657 |
| Casein kinase II subunit beta OS=Schistocephalus solidus OX=70667 GN=SSLN_LOCUS14489 PE=3 SV=1 | 1.95919047 | -0.253298 | -0.7343788 | 0.26926865 |
| SCP domain-containing protein OS=Schistocephalus solidus OX=70667 GN=SSLN_LOCUS193 PE=4 SV=1 | 1.47819531 | 0.83235832 | -1.0233186 | 0.49255657 |
| Uncharacterized protein OS=Schistocephalus solidus OX=70667 GN=SSLN_LOCUS5496 PE=4 SV=1 | 1.47819531 | 0.83235832 | -1.0233186 | 0.49255657 |
| UCR_hinge domain-containing protein OS=Schistocephalus solidus OX=70667 GN=SSLN_LOCUS11346 PE=3 SV=1 | 1.47819531 | 0.83235832 | -1.0233186 | 0.49255657 |
| Polypeptide N-acetylgalactosaminyltransferase OS=Schistocephalus solidus OX=70667 GN=GALT6 PE=3 SV=1 | 1.95919047 | -0.253298 | -0.7343788 | 0.26926865 |
| ATE_C domain-containing protein OS=Schistocephalus solidus OX=70667 GN=SSLN_LOCUS1820 PE=4 SV=1 | 1.47819531 | 0.83235832 | -1.0233186 | 0.49255657 |
| Plug_translocon domain-containing protein OS=Schistocephalus solidus OX=70667 GN=SSLN_LOCUS2320 PE=3 SV=1 | 1.95919047 | -0.253298 | -0.7343788 | 0.26926865 |
| Uncharacterized protein OS=Schistocephalus solidus OX=70667 GN=SSLN_LOCUS2429 PE=4 SV=1 | 1.47819531 | 0.83235832 | -1.0233186 | 0.49255657 |
| Uncharacterized protein OS=Schistocephalus solidus OX=70667 GN=SSLN_LOCUS7290 PE=4 SV=1 | 1.47819531 | 0.83235832 | -1.0233186 | 0.49255657 |
| Sec16_C domain-containing protein OS=Schistocephalus solidus OX=70667 GN=SSLN_LOCUS8651 PE=3 SV=1 | 1.91981946 | -0.335704 | -0.4974628 | 0.27466688 |
| Uncharacterized protein OS=Schistocephalus solidus OX=70667 GN=SSLN_LOCUS8725 PE=4 SV=1 | 1.47819531 | 0.83235832 | -1.0233186 | 0.49255657 |
| ANK_REP_REGION domain-containing protein OS=Schistocephalus solidus OX=70667 GN=SSLN_LOCUS9543 PE=4 SV=1 | 1.47819531 | 0.83235832 | -1.0233186 | 0.49255657 |
| Nuclear pore complex protein Nup98-Nup96 OS=Schistocephalus solidus OX=70667 GN=SSLN_LOCUS9851 PE=3 SV=1 | 1.91981946 | -0.335704 | -0.4974628 | 0.27466688 |
| Uncharacterized protein OS=Schistocephalus solidus OX=70667 GN=SSLN_LOCUS9860 PE=3 SV=1 | 1.95151089 | -0.2845542 | -0.5914534 | 0.12218842 |
| Uncharacterized protein OS=Schistocephalus solidus OX=70667 GN=SSLN_LOCUS10362 PE=4 SV=1 | 1.47819531 | 0.83235832 | -1.0233186 | 0.49255657 |
| Uncharacterized protein OS=Schistocephalus solidus OX=70667 GN=SSLN_LOCUS10761 PE=4 SV=1 | 1.47819531 | 0.83235832 | -1.0233186 | 0.49255657 |
| Uncharacterized protein OS=Schistocephalus solidus OX=70667 GN=SSLN_LOCUS13095 PE=4 SV=1 | 1.47819531 | 0.83235832 | -1.0233186 | 0.49255657 |
| Uncharacterized protein OS=Schistocephalus solidus OX=70667 GN=SSLN_LOCUS13671 PE=4 SV=1 | 1.47819531 | 0.83235832 | -1.0233186 | 0.49255657 |
| Uncharacterized protein OS=Schistocephalus solidus OX=70667 GN=SSLN_LOCUS14793 PE=4 SV=1 | 1.95919047 | -0.253298 | -0.7343788 | 0.26926865 |
| Tudor domain-containing protein OS=Schistocephalus solidus OX=70667 GN=SSLN_LOCUS16527 PE=4 SV=1 | 1.95919047 | -0.253298 | -0.7343788 | 0.26926865 |
| LSM14 domain-containing protein OS=Schistocephalus solidus OX=70667 GN=SSLN_LOCUS18830 PE=4 SV=1 | 1.95919047 | -0.253298 | -0.7343788 | 0.26926865 |
| Nudix hydrolase domain-containing protein OS=Schistocephalus solidus OX=70667 GN=SSLN_LOCUS88 PE=4 SV=1 | 1.47819531 | 0.83235832 | -1.0233186 | 0.49255657 |
| Uncharacterized protein OS=Schistocephalus solidus OX=70667 GN=SSLN_LOCUS5654 PE=4 SV=1 | 1.95919047 | -0.253298 | -0.7343788 | 0.26926865 |
| Palmitoyltransferase OS=Schistocephalus solidus OX=70667 GN=SSLN_LOCUS2951 PE=3 SV=1 | 1.47819531 | 0.83235832 | -1.0233186 | 0.49255657 |
| Uncharacterized protein OS=Schistocephalus solidus OX=70667 GN=SSLN_LOCUS12875 PE=4 SV=1 | 1.95919047 | -0.253298 | -0.7343788 | 0.26926865 |
| Uncharacterized protein OS=Schistocephalus solidus OX=70667 GN=SSLN_LOCUS9518 PE=4 SV=1 | 1.95919047 | -0.253298 | -0.7343788 | 0.26926865 |
| SH3 domain-containing protein OS=Schistocephalus solidus OX=70667 GN=SSLN_LOCUS288 PE=4 SV=1 | 1.47819531 | 0.83235832 | -1.0233186 | 0.49255657 |
| Uncharacterized protein OS=Schistocephalus solidus OX=70667 GN=SSLN_LOCUS16326 PE=3 SV=1 | 1.95151089 | -0.2845542 | -0.5914534 | 0.12218842 |

Supplementary Table S4: MANOVA

| **Comparison Type** | **Df** | **Pillai’s Trace** | **Approximate F-statistic** | **Num Df** | **Den Df** | **Pr(>F)** | **K** |
| --- | --- | --- | --- | --- | --- | --- | --- |
| ESP vs Tissue (among lake populations) | 1 | 0.87337 | 172.42 | 2 | 50 | <2.2e-16 | 2 |
| ESP only (among lake populations) | 3 | 1.1893 | 3.4482 | 12 | 63 | .00063111 | 4 |
| Tissue only (among lake populations) | 3 | 1.6342 | 6.5808 | 12 | 66 | 1.444e-07 | 4 |
| Tissue+Lake population difference | 1 | .94215 | 158.784 | 4 | 39 | <2.2e-16 | 8 |
| Lake population difference | 3 | .90875 | 4.454 | 12 | 123 | 6.62e-06 | 8 |
| (Tissue*Population)+Mass | 1 | .18228 | 2.173 | 4 | 39 | .090006 | 8 |
| Tissue:Genotype | 3 | .67442 | 2.973 | 12 | 123 | .001135 | 8 |
